# Cytoplasmic tail of the putative polycystin channel Pkd2 regulates its clustering in the fission yeast eisosomes

**DOI:** 10.1101/2022.08.11.503631

**Authors:** Mamata Malla, Debatrayee Sinha, Pritha Chowdhury, Benjamin Thomas Bisesi, Qian Chen

## Abstract

Polycystins are a family of conserved ion channels, mutations of which lead to human genetic disorder Autosomal Dominant Polycystic Kidney Disease. The unicellular model organism fission yeast *Schizosacchromyces pombe* possesses single essential polycystin Pkd2 that localizes to the plasma membrane and is required for cell proliferation. Here, we carried out a functional analysis of Pkd2 based on its Alphafold predicted structure. It consisted of N-terminal lipid-binding (LBD), central transmembrane (TMD) and C-terminal cytoplasmic (CCD) domains. LBD assumes a unique immunoglobulin-fold, while TMD contains nine transmembrane helices. Both were essential. Although the mostly disordered CCD was not, its removal led to clustering of Pkd2 in eisosomes, a microdomain of the plasma membrane. Inhibiting eisosome assembly prevented the clustering but disrupting ER-PM contacts further increased it. Pkd2 shared similar structure with two other putative channels Trp663 and Trp1322, but their intracellular localization and function diverged from each other. Replacing LBD with that of Trp663 partially restored the function of Pkd2, but TMD could not be replaced by either that of Trp663 or human polycystins. We concluded that both the plasma membrane microdomains and cytoplasmic tail of Pkd2 regulate the cell surface clustering of this putative ion channel.

## Introduction

Polycystins are a family of transmembrane proteins that are evolutionarily conserved from yeast to humans. Loss of function mutations in either one of two human polycystin genes, which encode PC1 and PC2 respectively, result in one of the most common genetic disorders Autosomal Dominant Polycystic Kidney Disease (ADPKD) (Wu et al., 1998). Of the two human polycystins, PC1 is a putative mechanosensor localized on the plasma membrane as well as the primary cilium (Geng et al., 1996; Parnell et al., 2018; Su et al., 2015). In comparison, PC2 oligomerizes into a calcium-permissive transient receptor potential (TRP) channel that mostly localizes on the endoplasmic reticulum (ER) (Foggensteiner et al., 2000) in addition to the primary cilium (Liu et al., 2018; Pazour et al., 2002). These two human polycystins are also capable of assembling into a hetero-tetrameric channel (Grieben et al., 2017; Su et al., 2018). Nevertheless, their cellular functions remain unclear.

The fission yeast *Schizosacchromyces pombe* possesses just one essential polycystin homologue Pkd2. This putative TRP channel (Palmer et al., 2005) localizes to the plasma membrane (PM) in a cell cycle-dependent manner, concentrating at the cell tips during interphase growth and translocating to the equatorial plane during cell division (Morris et al., 2019). Such dynamic localization of Pkd2 depends on the actin cytoskeleton as well as intracellular membrane trafficking.

The function of Pkd2 is essential for cell proliferation including both cell division and cell growth (Morris et al., 2019; Sinha et al., 2022). In cytokinesis, the last stage of cell division, Pkd2 modulates the contractile ring constriction at the beginning and promotes cell separation at the very end (Morris et al., 2019). Its activity also antagonizes that of the essential yeast Hippo-like signaling pathway Septation Initiation Network (SIN). As a result, mutations of *pkd2* partially rescue the lethal SIN mutations (Sinha et al., 2022). During cell growth, Pkd2 promotes the tip extension by regulating the turgor pressure. A temperature-sensitive mutation *pkd2-B42* leads to significantly reduced cell tip growth and a failure to maintain the cell volume homeostasis, a unique phenotype termed “Deflation” (Sinha et al., 2022). In contrast to Pkd2, two other putative fission yeast TRP channels Trp663 and Trp1322 are not essential. Their cellular function and intracellular localization are unknown (Morris et al., 2019).

In this study, we took advantage of the newly available Alphafold predicted structure of Pkd2 to examine its structure-function relationship. Pkd2 is projected to possesses three unique domains: LBD, TMD and CCD. Based on this prediction, we constructed a series of Pkd2 truncation mutants, characterized their localization by fluorescence microscopy, and tested their function through genetic analyses. Both LBD and TMD are essential for the plasma membrane localization and function of Pkd2. Although CCD was not essential for the function, its removal led to clustering of Pkd2 on the plasma membrane, in eisosomes. Inhibition of eisosome assembly prevented the clustering, while disruption of the ER-PM contacts substantially increased it. We also compared the domains of Pkd2 with those of Trp663, Trp1322, a Pkd2 homologue in *S. japonicus* and human polycystins by constructing various chimeras. Our results showed that both LBD and TMD of Pkd2 are conserved in fission yeast species.

## Results

### Alphafold predicted tertiary structure of Pkd2

Since the structure of fission yeast Pkd2 has not been experimentally determined, we employed Alphafold to predict the putative domains. Preliminary analysis of the primary and secondary structures of Pkd2 suggested that it possesses a 23a.a. (amino acid) N-terminal signal peptide and three likely domains, the 147a.a. putative lipid-binding domain (LBD), followed by the 405a.a. transmembrane domain (TMD) and the 135a.a. C-terminal cytoplasmic domain (CCD) (Fig. 1A). Largely agreeing with this, Alphafold (Jumper et al., 2021) predicted a tertiary structure of three distinct domains (Fig. 1B), with high confidence on LBD and TMD. In this putative structure, the extracellular N-terminal LBD adopts an immunoglobulin (Ig)-like fold (Bork et al., 1994), consisting of two opposing β-sheets, each with four anti-parallel β-strands (A to H) (Fig. 1C). A cavity is projected to form between the two β-sheets. Like many Ig-like domain, LBD possessed pairing cysteine residues, four in total at the positions of 36, 92, 100 and 155 respectively. They are projected to form two pairs, one between the 1^st^ and 4^th^ cysteines and the other between 2^nd^ and 3^rd^ (Fig. 1C), suggesting the existence of two stabilizing disulfide bridges in LBD. Unlike most polycystins, TMD of Pkd2 is projected to contain nine transmembrane helices (Fig. 1D). It is predicted to have extensive contacts with LBD. CCD, whose structure was predicted with the lowest confidence among three domains, consists of mostly disordered sequences in addition to a short coiled-coil motif (Fig. 1B). Based on these computational analyses, we concluded that Pkd2 most likely consists of three structurally distinct domains of LBD, TMD and CCD.

**Figure 1.**
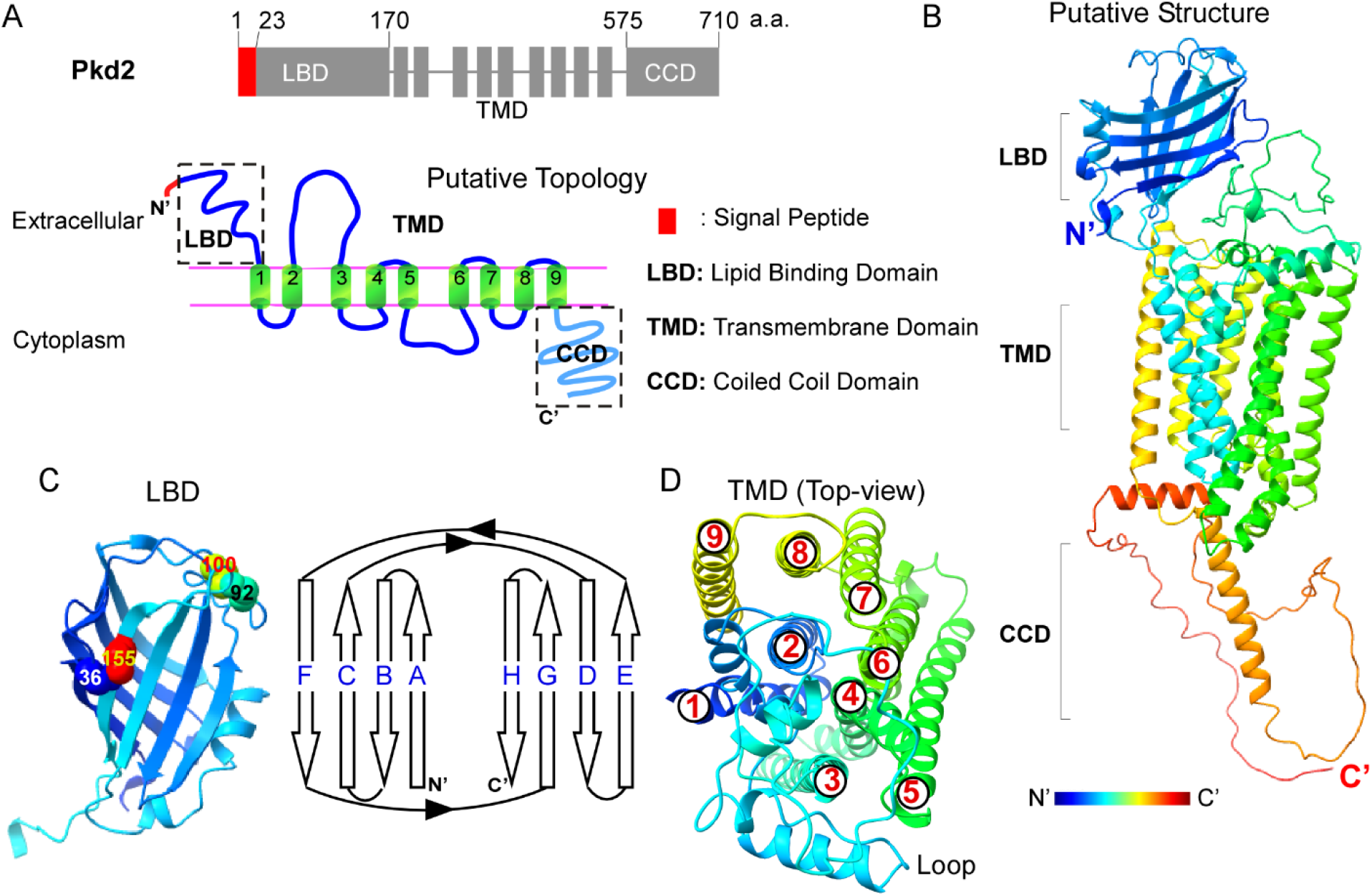
Alphafold predicted structure of Pkd2. **(A)** Schematics of the primary structure (top) and topology (bottom) of Pkd2. The topology was predicted using Protter. **(B)** Ribbon diagram of the Alphafold predicted tertiary structure (rainbow colored) of Pkd2. The signal peptide has been removed. The structure was rendered using ChimeraX (UCSF) **(C-D)** Ribbon diagram of the structures of LBD (C) and TMD (D). (C) Left: side-view of LBD with two pairs of cysteines labelled and rendered with surface-view. Right: 2D topology of 8 anti-parallel β-strands (A to H) in LBD. (D) Top view of the 9 transmembrane helices (1 to 9) in TMD.

### Both LBD and TMD of Pkd2 are required for its plasma membrane localization and function

We expressed each of the three Pkd2 domains separately to determine their localization and function. None of the ectopically expressed LBD-GFP, CCD-GFP and TMD-GFP (see Materials and Methods) localized to the plasma membrane. The former two diffused throughout the cytoplasm (Fig. S1A and S1C), while last one localized instead at the cortical ER, marked by mCherry-ADEL (Fig. S1B). Since TMD is projected to possess more transmembrane helices than the six in a canonical TRP channel, we systematically removed one or more of the nine helices at a time to generate a series of truncation mutants (Table S1). Again, all of these mutants failed to reach the plasma membrane (Table S1). Functionally, none of these close to a dozen Pkd2 truncation mutants rescued the temperature-sensitive mutant *pkd2-B42* (Sinha et al., 2022) (Fig. S1D-E and Table S1). Thus, none of the three domains is sufficient on its own for either the plasma membrane localization or function of Pkd2.

Next, we determined which one of these three domains is absolutely required for the function of Pkd2. Since *pkd2* is an essential gene for fission yeast viability (Morris et al., 2019), we tested this by replacing the full-length gene with three truncation mutants *pkd2Δ LBD, pkd2Δ TMDΔ CCD* or *pkd2Δ CCD* respectively at the endogenous *pkd2* locus. The first two mutants were inviable (Fig. 2A and 2B). In contrast, the last one without CCD was viable and its growth was similar to the wild-type at temperatures ranging from 25°C to 36°C (Fig. 2C). We concluded that both LBD and TMD domains are essential for the function of Pkd2.

**Figure 2.**
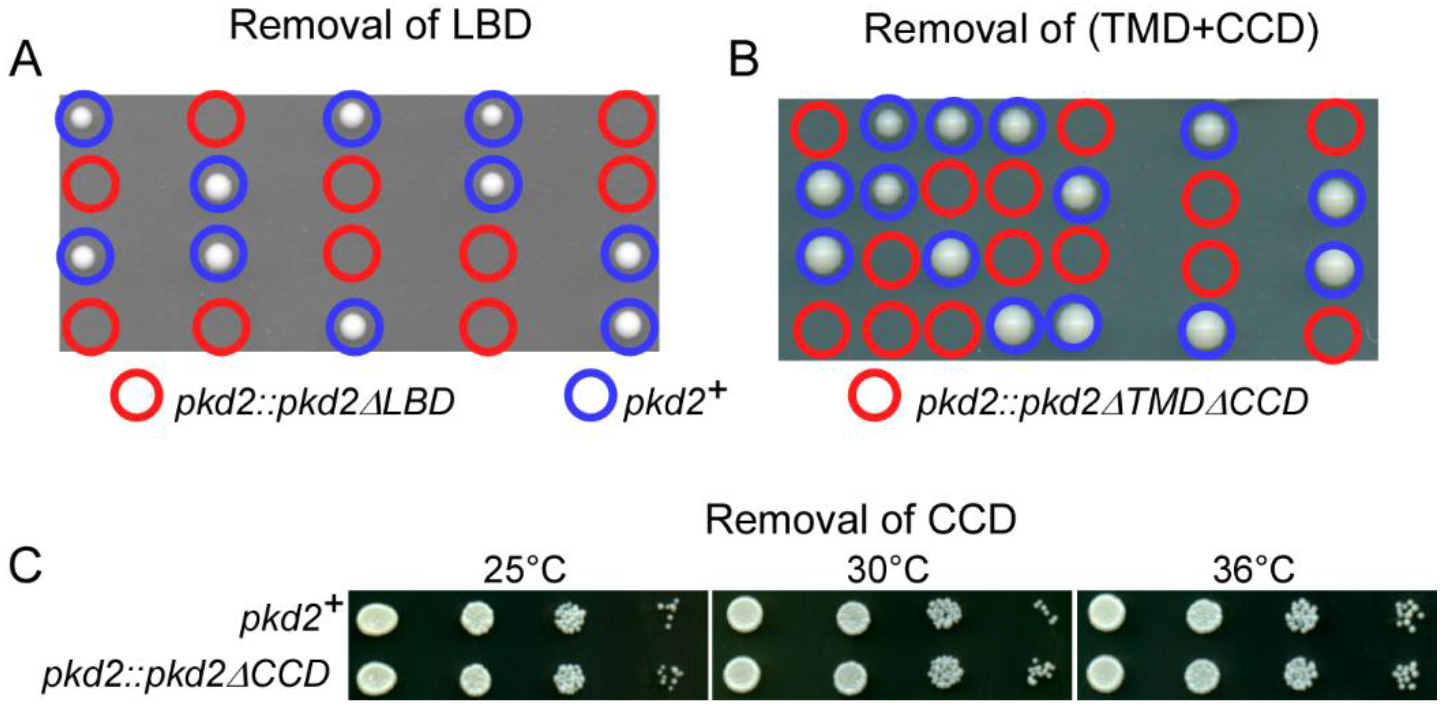
Both LBD and TMD of Pkd2 are essential for its function. **(A-B)** Tetrad-dissection plates of sporulated *pkd2*^*+*^/*pkd2ΔLBD* (A) or *pkd2*^*+*^/*pkd2ΔTMDΔCCD* (B). Both *pkd2::pkd2Δ LBD* (red circle) and *pkd2::pkd2Δ TMDΔ CCD* (red circle) were inviable. **(C)** Ten-fold dilution series of yeast on YE5s plates. *pkd2:pkd2Δ CCD* grew similarly to the wild-type.

### CCD-less Pkd2 clustered in eisosomes on the cell surface

To determine the potential role of CCD, we first characterized the intracellular localization of Pkd2Δ CCD using fluorescence microscopy. This truncation mutant, when tagged with GFP, continued to localize to the plasma membrane, either at the cell tip during cell growth and division as well or at the equatorial division plane during cytokinesis (Fig. 3A). However, unlike the full-length protein which was found in many intracellular membrane structures (Morris et al., 2019), this truncated Pkd2 mostly remained free from them (Fig. 3B). Most surprisingly, Pkd2Δ CCD-GFP clustered throughout the cell surface, forming a network of short filaments (Fig. 3A). Therefore, CCD does have a critical role in modulating the internalization of Pkd2 and its clustering.

**Figure 3.**
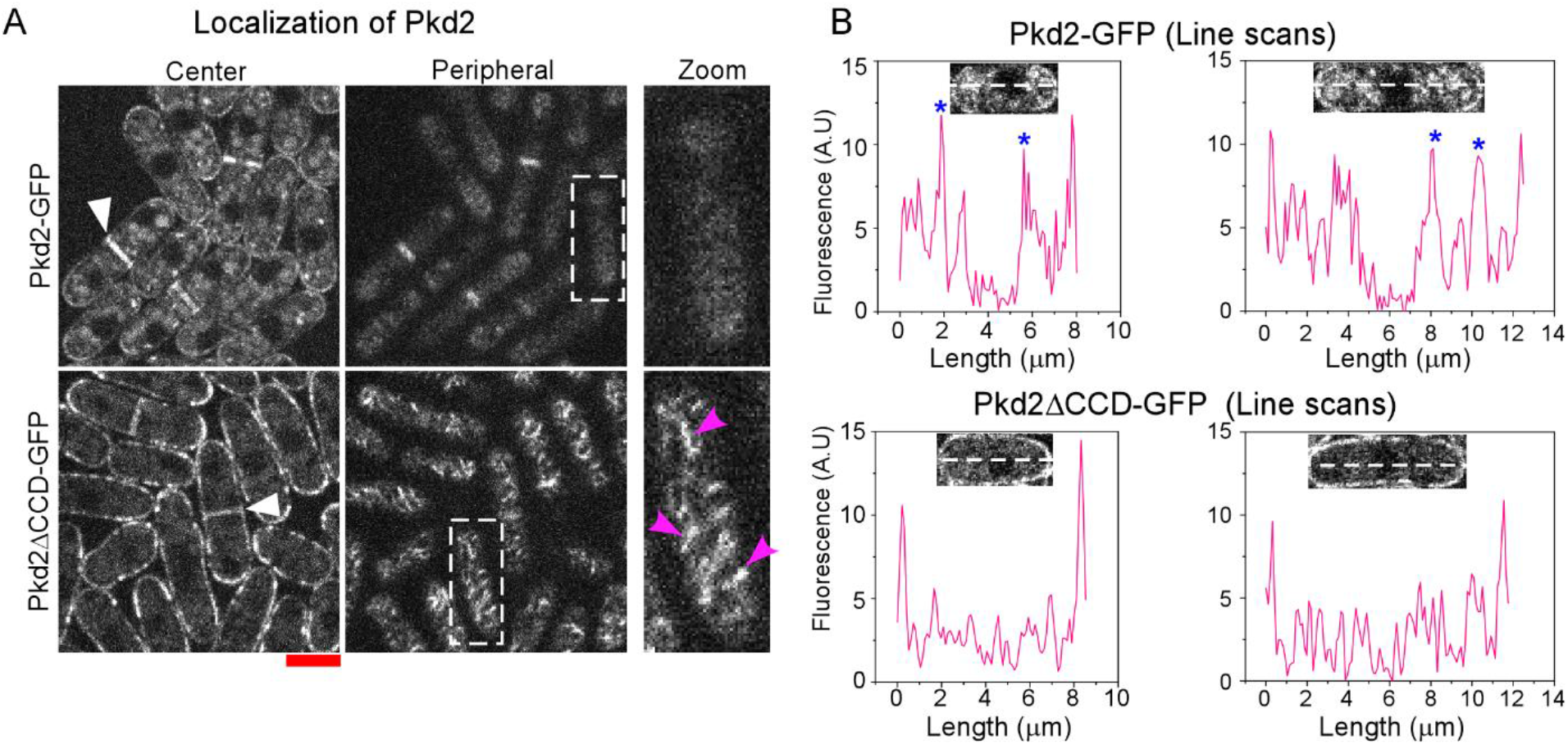
Removal of CCD reduces internalization of Pkd2 from the plasma membrane. **(A)** Micrographs of the cells expressing either Pkd2-GFP (top) or Pkd2ΔCCD-GFP (bottom). Both the center (left) and the peripheral (middle) slices of the z-series are shown. A magnified view of the surface of representative cells (dashed line) is shown on the right. White arrowhead: equatorial plane. Purple arrowhead: the filaments on cell surface where Pkd2ΔCCD-GFP clustered. **(B)** Fluorescence intensity plot of the line-scans of either Pkd2-GFP (top) or Pkd2ΔCCD-GFP (bottom) expressing cells (insert). The center slice of the respective Z-series was used.

To determine the nature of these novel Pkd2Δ CCD filaments, we compared them to the well-known microdomain of the plasma membrane, eisosomes. As found previously (Kabeche et al., 2011), the fission yeast eisosomes, marked by an essential component Pil1-mCherry, distributed as a filamentous network on the cell surface (Fig. 4A). These filaments were largely absent from the cell tips and the equatorial division plane. In comparison, the filaments of Pkd2Δ CCD-GFP were shorter, and they distributed far more evenly throughout the cell surface (Fig. 4A). Nevertheless, in the mid-portion of non-dividing cells, these two types of filaments often partially overlapped with each other (Fig. 4A). The average Pearson’s co-localization coefficient between Pkd2Δ CCD and Pil1 on the cell surface was μ0.3 (Fig. 4B). In comparison, the full-length Pkd2 did not cluster in eisosomes (Fig. 4A and 4B). It did not co-localize with Pil1 at all, with a Pearson’s coefficient that was close to zero. We concluded that without CCD Pkd2 largely clusters along the filaments of eisosomes on the cell surface.

**Figure 4.**
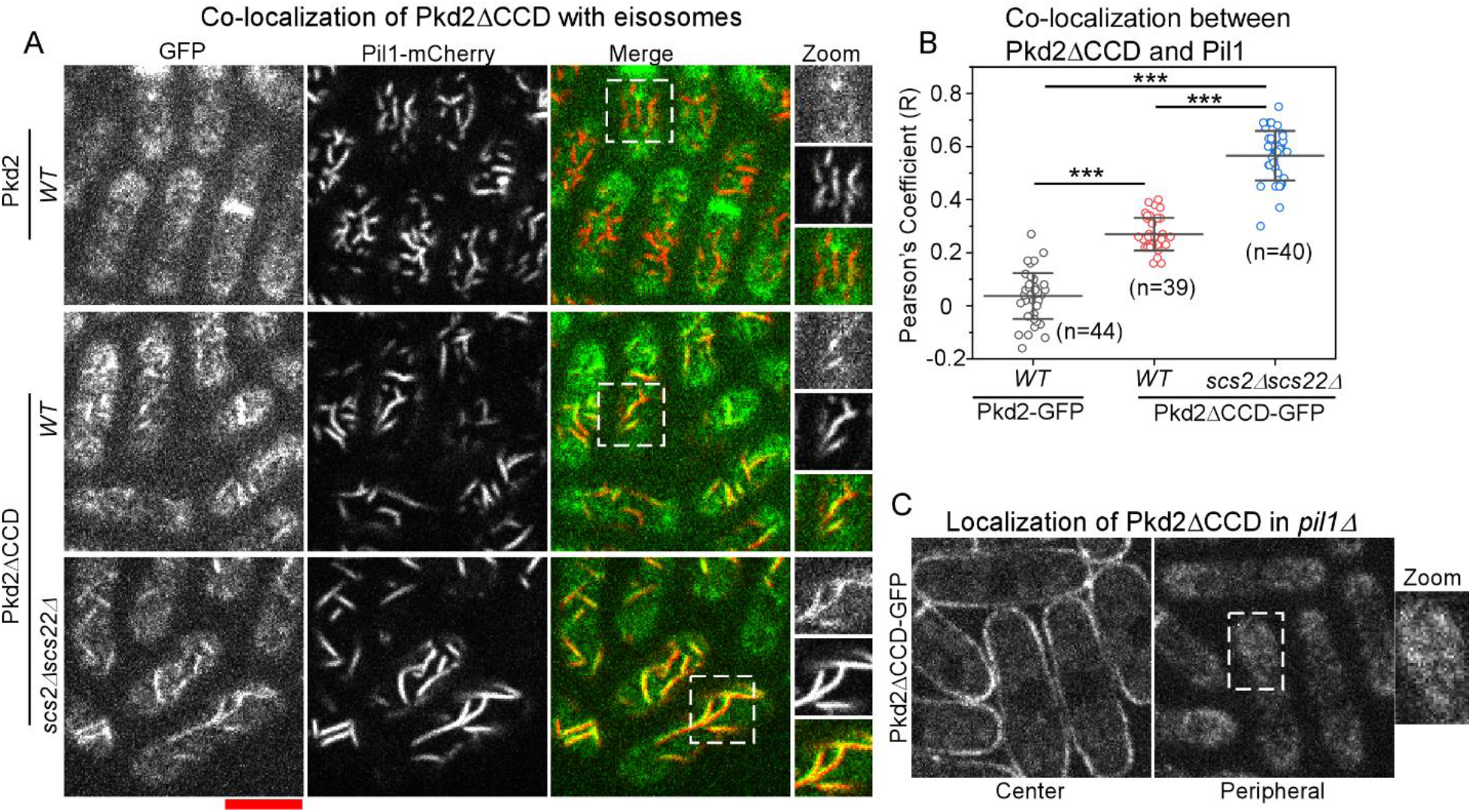
Pkd2Δ CCD clusters in eisosomes on the cell surface. **(A)** Micrographs of the cells expressing the eisosome marker Pil1-mCherry (red in the merged view) and either Pkd2-GFP (top, green in merged view) or Pkd2ΔCCD-GFP (middle and bottom). *WT*: wild-type. Single planes of the cell surface are shown. Right: magnified views of the cell surface (dashed line). **(B)** Plot of the average Pearson’s co-localization coefficient between Pil1-mCherry and either Pkd2-GFP or Pkd2ΔCCD-GFP in cells. Scale: 5μm. ***: P≤0.0001 (Two-tailed student’s t-test). Number: total number of cells analyzed. Data is from at least two independent biological repeats. **(C)** Micrograph of the *pil1*Δ cells expressing Pkd2ΔCCD-GFP. Left: center slice. Right: cell surface view, magnified view of the cell surface is shown in zoom (dashed line).

Next, we determined whether eisosomes are required for the cell surface clustering of Pkd2Δ CCD. Since Pil1 is essential for the eisosome assembly (Kabeche et al., 2011), we examined the localization of Pkd2Δ CCD-GFP in the *pil1Δ* mutant. Indeed, clustering of Pkd2Δ CCD on the cell surface disappeared completely in the mutant cells (Fig. 4C). All the Pkd2Δ CCD-marked filaments, including those at the cell tips and the division plane, dissipated (Fig. 4C). As a result, Pkd2Δ CCD-GFP distributed homogenously on the plasma membrane (Fig. 4C). We concluded that eisosomes promote the clustering of Pkd2 on the cell surface.

As the fission yeast eisosomes are closely associated with the ER-PM contacts (Ng et al., 2020), we asked whether these membrane contact sites regulate the clustering of Pkd2Δ CCD. We characterized the localization of Pkd2Δ CCD-GFP in the mutant of Scs2 and Scs22, two VAP (VAMP-associated protein) that are tethers between the ER and PM (Manford et al., 2012; Zhang et al., 2012). As reported previously (Ng et al., 2020), the number of the eisosome filaments decreased in *scs2Δ scs22Δ* cells, compared to the wild type (Fig. 4A). However, we also found that the average length of these eisosome filaments increased, and they distributed more broadly throughout the cell surface including the tips, compared to those in the wild-type cells (Fig. 4A). Surprisingly, in these *scs2Δ scs22Δ* cells, clustering of Pkd2Δ CCD increased significantly (Fig. 4A). Those Pkd2Δ CCD-marked filaments became longer and more distinct.

They almost always overlapped with the eisosome filaments. Indeed, the Pearson’s co-localization coefficient between Pkd2Δ CCD and Pil1 doubled to 0.6 in *scs2Δ scs22Δ* cells, compared to the wild-type cells (Fig 4B). In contrast, full-length Pkd2 did not cluster in eisosomes at all even in these *scs2Δ scs22Δ* cells (Fig. S2B). We concluded that ER-PM contacts modulate the clustering of Pkd2Δ CCD in eisosomes on the cell surface.

Finally, we examined how the clustering of Pkd2Δ CCD affects its translocation to the equatorial division plane. As fission yeast cells enter cytokinesis, the full-length protein moves to the equatorial plane, coincided with the start of the contractile ring constriction (Fig. 5A and 5B). Pkd2Δ CCD-GFP did the same (Fig. 5B). And just like the full-length Pkd2, the molecular number of Pkd2Δ CCD peaked as the ring constriction completed (Fig. 5C). However, compared to the full-length protein, the number of the Pkd2Δ CCD molecules at the division plane decreased by μ50% (Fig. 5C), despite their similar expression levels (Fig. S2A). More difference lied in their kinetics. While the number of Pkd2Δ CCD molecules remained constant throughout the cell separation, the number of the full-length protein decreased gradually (Fig. 5C). Thus, CCD promotes the translocation of Pkd2 to the equatorial plane during cytokinesis.

**Figure 5.**
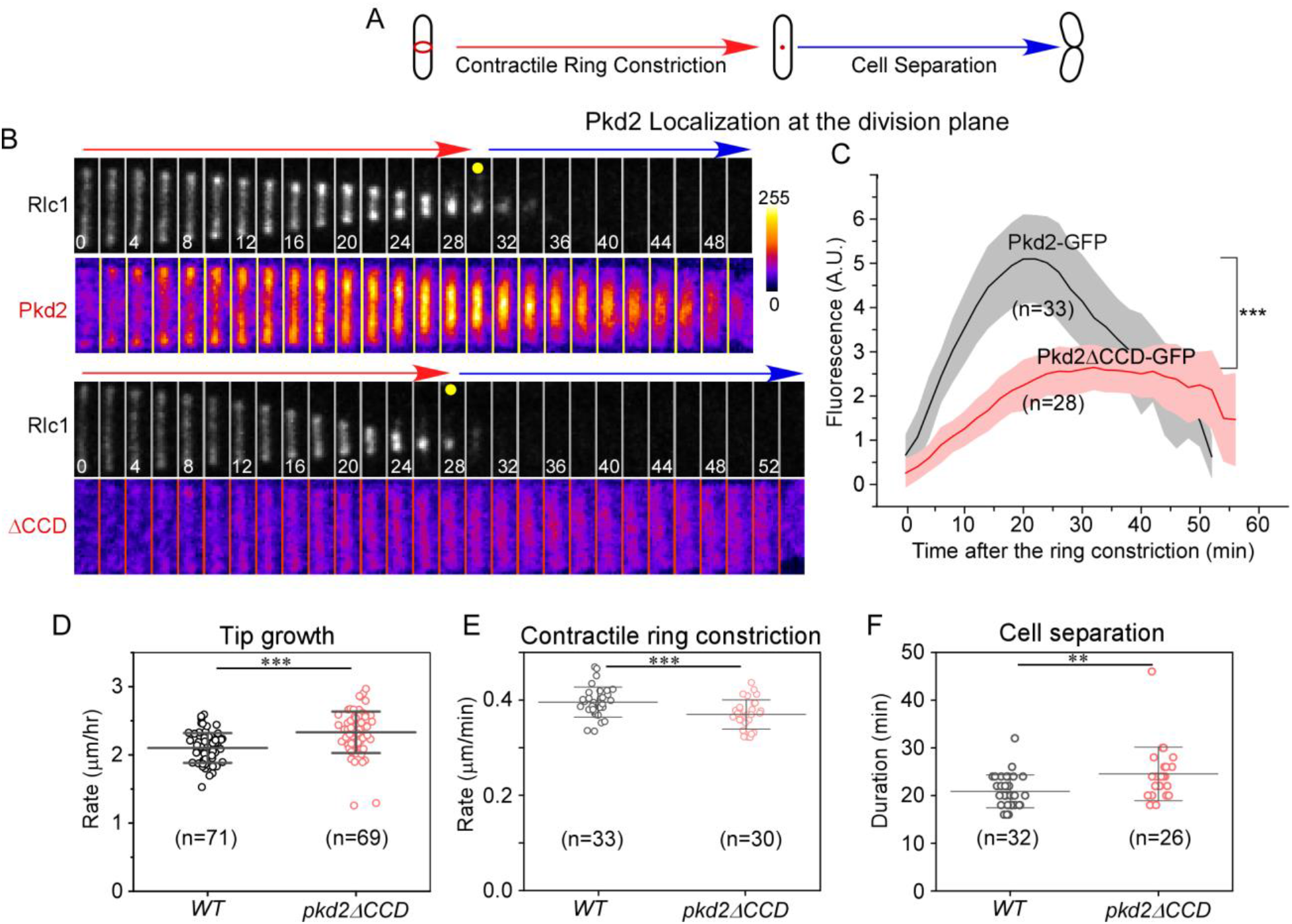
Pkd2Δ CCD remains largely functional during both cell growth and cytokinesis. **(A-C)** Localization of Pkd2-GFP at the equatorial plane during cytokinesis. (A) A schematic of the fission yeast cytokinesis. Red: actomyosin contractile ring. (B) Time-series showing the division plane of a cell expressing either Pkd2-GFP (pseudo-colored, top) or Pkd2ΔCCD-GFP (bottom) and the contractile ring marker Rlc1-tdTomato (gray). Numbers: time in minutes after the start of the contractile ring constriction. Bar: fluorescence intensity scale. (C) Time-course of the fluorescence intensities of Pkd2-GFP and Pkd2ΔCCD-GFP in the division plane. **(D)** Tip growth rates of wild-type (*WT)* and *pkd2ΔCCD* mutant cells. **(E-F)** Progression of cytokinesis in *pkd2Δ CCD* mutant cells is normal. Dot plots of the contractile ring constriction rate (D) and the duration of cell separation (E). Bars represent average ± standard deviation. **: P ≤0.001, ***: P≤0.0001 (Two-tailed student’s t-test). Data is pooled from at least two independent biological repeats.

While it is clear that CCD regulates the localization of Pkd2, we still wondered whether it has any non-essential role in the function of this transmembrane protein. We thus quantified the cell tip growth and cytokinesis of *pkd2::pkd2Δ CCD* mutant through time-lapse microscopy. In comparison to the wild-type cells, *pkd2::pkd2Δ CCD* cells extended their tip at a similar rate, only μ10% higher (Fig. 5D). Likewise, both the contractile ring constriction and cell separation proceeded mostly unperturbed in the truncation mutant cells (Fig. 5E and 5F). We concluded that CCD is largely dispensable for the cellular functions of Pkd2 despite its role in regulating the clustering.

### A chimera of Trp663 and Pkd2 is partially functional

Surprisingly, Pkd2 is not alone in its structure among the fission yeast TRP channels. Alphafold predicted that two others, Trp663 and Trp1322 (Lamas et al., 2020; Morris et al., 2019) shared a similar domain structure (Fig. 6A), despite the overall sequence identities among these three proteins is just μ20%. All three possess a similar extracellular LBD, that shared the unique Ig-like fold consisted of eight anti-parallel β-strands (Fig. 6A). All of their putative TMD structure contain 9 transmembrane helices, despite that the secondary structure analysis (Protter) predicted 8 and 11 for Trp663 and Trp1322 respectively (Omasits et al., 2014). We concluded that all three putative TRP channels of fission yeast possess similar tertiary structures especially in their LBDs and TMDs.

**Figure 6.**
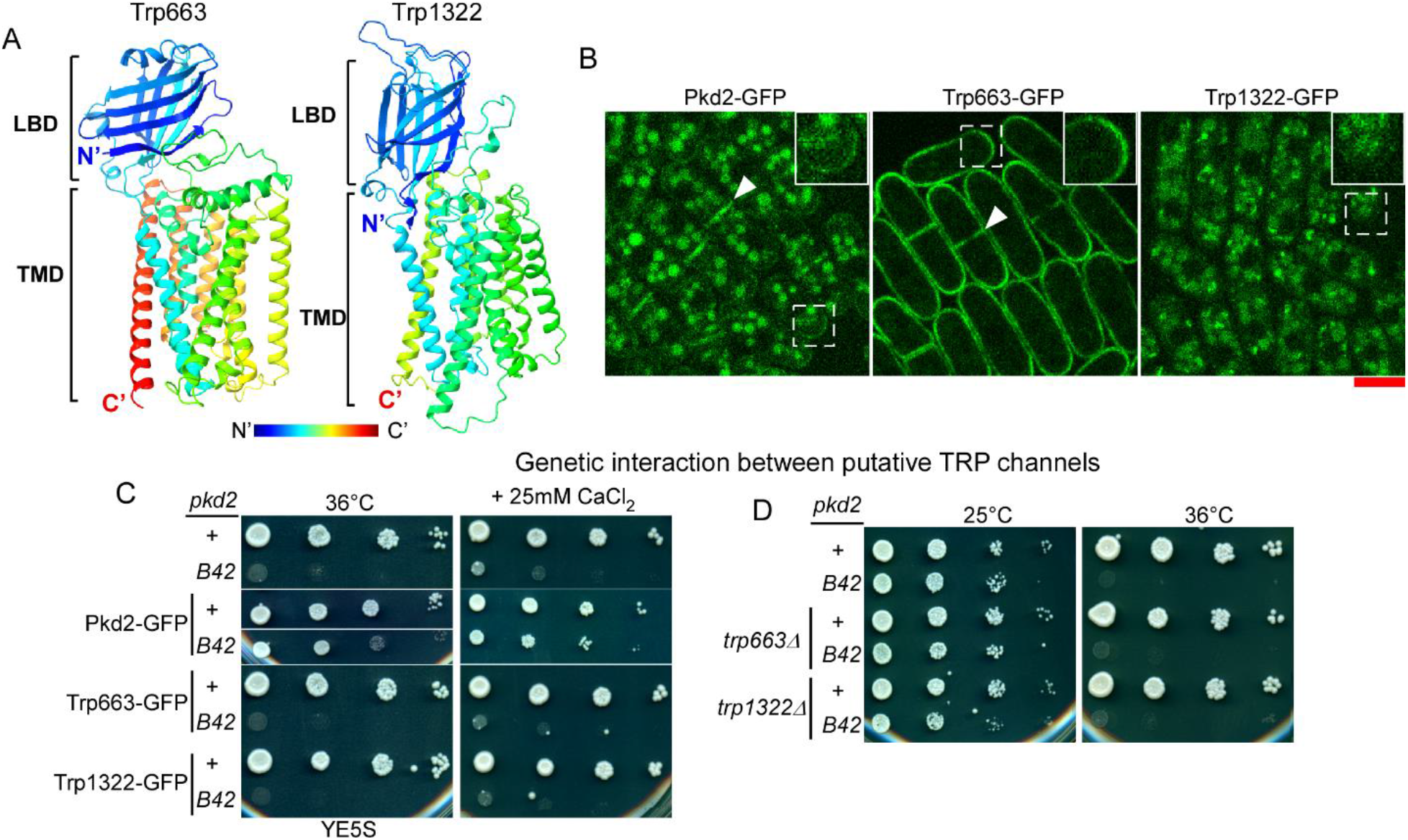
The structures of Trp663 and Trp1322 are similar to that of Pkd2, but their localization and function differ. **(A)** Ribbon diagram of the Alphafold predicted structures of Trp663 and Trp1322 (rainbow colored). Their N-terminal signal peptide and mostly disordered C-terminal cytoplasmic tail are not shown. **(B)** Micrographs of cells ectopically expressing either Pkd2-GFP (left), or Trp663-GFP (middle) or Trp1322-GFP (right). Arrowhead: the equatorial division plane. Magnified views show the difference in localization on the plasma membrane at cell tips (dashed lines) Scale: 5μm **(C-D)** Genetic interaction between *pkd2, trp663* and *trp1322*. Ten-fold dilution series of yeast. (C) The wild-type (*pkd2*^*+*^) or *pkd2-B42* ectopically expressing either Pkd2, Trp663 or Trp1322. (D) Double mutants of *pkd2-B42* and *trp663Δ* or *trp1322Δ*.

We were then curious whether these three TRP channels distribute similarly in cells. Unlike Pkd2, the intracellular localization of endogenous Trp663 and Trp1322 remained unknown, because both expressed at a relatively low level (Morris et al., 2019). Therefore, we expressed all three ectopically under the control of a strong promoter. As a control, ectopically expressed Pkd2-GFP largely maintained the endogenous localization (Fig. 6B) with slightly increased distribution towards the intracellular membrane structures. In comparison, Trp663-GFP exclusively localized to the plasma membrane (Fig. 6B). It distributed homogenously on the cell surface, exhibiting neither clustering nor enrichment at the growth zones of either cell tip or the division plane. Further, it stayed largely free from the intracellular membrane structures in which Pkd2 was often found. In contrast to Pkd2 and Trp663, Trp1322-GFP was completely absent from the plasma membrane (Fig. 6B). Instead, it mostly localized to the cytoplasmic puncta. We concluded that these three putative TRP channels indeed occupy different intracellular compartments despite their similar structures.

Next, we compared the cellular functions of these three channels. We determined whether there is any functional overlap between Pkd2 and either Trp663 or Trp1322 through genetic studies. As expected, over-expressed Pkd2 largely rescued *pkd2-B42* at the restrictive temperature of 36°C (Fig. S3A-C). In contrast, expression of either Trp663 or Trp1322 failed to rescue *pkd2-B42* (Fig 6C). Further, neither *trp663Δ* nor *trp1322Δ* exhibited any genetic interactions with *pkd2-B42* (Fig. 6D). We concluded that there is little overlap among the functions of three TRP channels.

Since both Pkd2 and Trp663 are present on the plasma membrane, we wondered whether their functions may have diverged due to their respective LBD or TMD domains. First, we constructed a chimera between these two by replacing LBD of Pkd2 with that of Trp663, Trp663^LBD^-Pkd2^(TMD-CCD)^ (Chimera I hereafter). This chimera, tagged with GFP, mostly stayed at the cortical ER (Fig 7A). Although it rescued the viability of *pkd2-B42* (Fig. 7C and Table S2), this chimera failed to revive *pkd2Δ* mutant, unlike the full-length protein (Fig. S4A and S4B). Secondly, we constructed another chimera by replacing both TMD and CCD of Pkd2 with those of Trp663, Pkd2^LBD^-Trp663^(TMD-CCD)^ (Chimera II). This chimeric protein tagged with GFP were found at both the cortical ER and the plasma membrane (Fig. 7B). However, it failed to even rescue *pkd2-B42* (Fig. 7C). We concluded that LBD is only partially conserved between Pkd2 and Trp663, but TMD is not.

**Figure 7.**
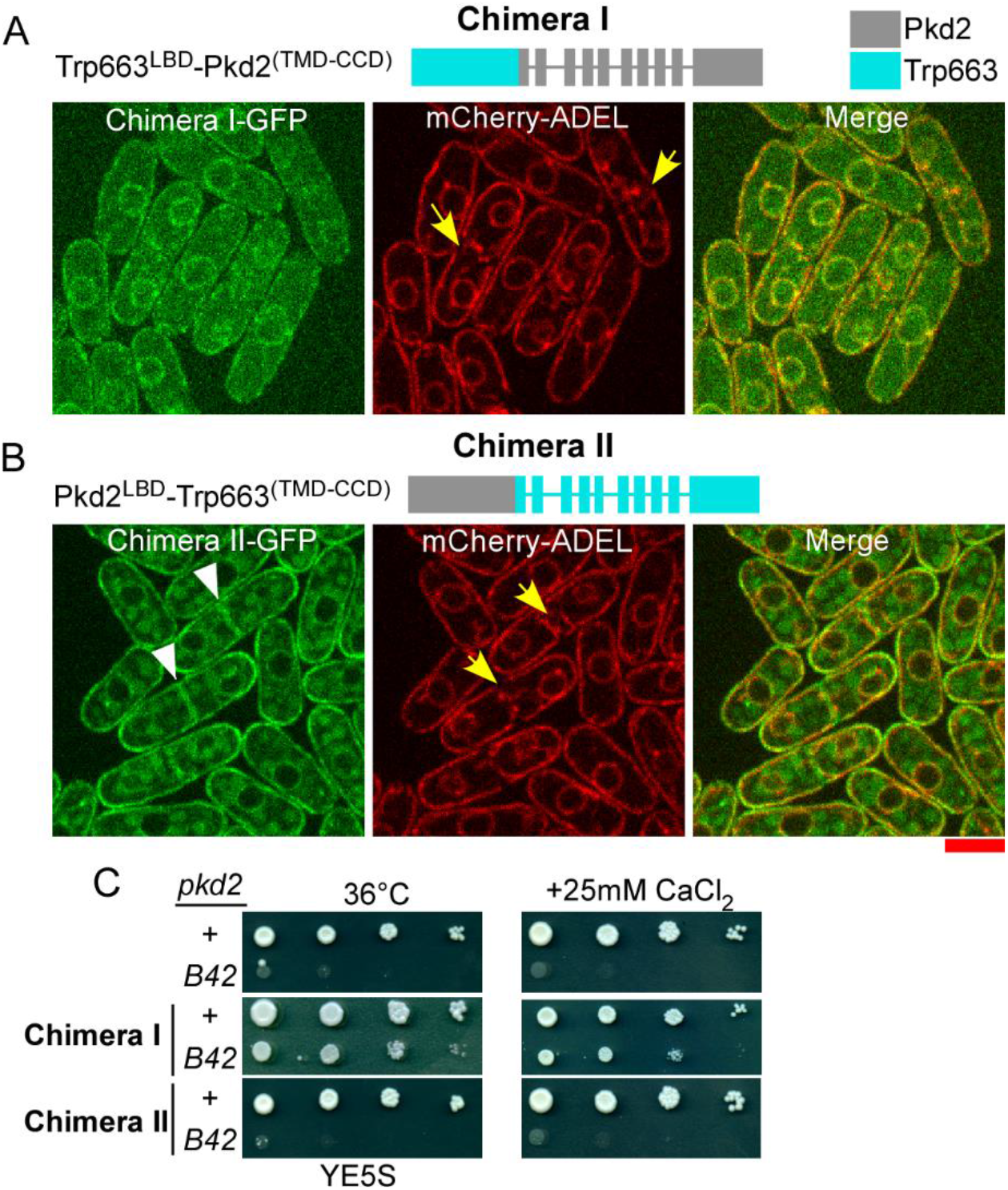
A chimera of Pkd2 and Trp663 rescues *pkd2-B42*. **(A)** Top: Schematic of Chimera I (Trp663^LBD^-Pkd2 ^(TMD-CCD)^) which replaced LBD of Pkd2 with that of Trp663. Bottom: micrographs of the cells expressing both Chimera I tagged with GFP (green) and the cortical ER marker mCherry-ADEL (red). **(B)** Top: Schematic of Chimera II (Pkd2^LBD^-Trp663^(TMD-CCD)^) which replaced TMD and CCD of Pkd2 with those of Trp663. Bottom: micrographs of the cells expressing both Chimera II tagged with GFP (green) and mCherry-ADEL (red). Arrow: the equatorial division plane. Scale: 5μm. **(C)** Ten-fold dilution series of yeast on either YE5S at 36°C (left) or YE5S plus 25mM CaCl2 (right) plates.

### Localization and function of Pkd2 are conserved among fission yeast species

To determine whether the structure of Pkd2 has been conserved in evolution, we first compared TMD to those of human polycystins. We constructed two chimeras between the yeast and human polycystins by replacing TMD with that of either human PC-1 or PC-2, Pkd2^LBD^-PC1^TMD^ (Chimera III) (Fig. S5A) and Pkd2^(LBD+TM1)^-PC2^TMD^ (Chimera IV) (Fig. S5B) respectively.

Whether expressed individually or together, both chimeras stayed at the cortical ER, absent from the plasma membrane (Fig. S5A-C). Most importantly, they failed to rescue *pkd2-B42* (Fig. S5D). We concluded that the TMD of fission yeast polycystin Pkd2 is not conserved in the human polycystins.

Next, we determined whether Pkd2 has been conserved in the other fission yeast species. Besides *S. pombe*, there are three other fission yeast species *S. octosporus (So), S. crypophilus (Sc)* and *S. japonicus (Sj)* (Rhind et al., 2011). Through BLAST search, we identified a single Pkd2 homologue in each of them. They share 63%, 62% and 49% sequence identities with Pkd2 respectively. For functional comparison, we chose *Sj*Pkd2 which is the most distant relative (Rhind et al., 2011). The putative topology of *Sj*Pkd2 was identical to that of Pkd2, as was its Alphafold-predicted structure (Fig. 8A). When ectopically expressed, *Sj*Pkd2-GFP localized to the plasma membrane, either at the cell tips during the growth or at the equatorial division plane during cytokinesis, as Pkd2 does (Fig. 8B). Importantly, *Sj*Pkd2-GFP rescued *pkd2-B42* and *pkd2Δ* mutants were viable by the ectopic expression of Pkd2(Fig. 8C-E). We concluded that the structure and function of the essential protein Pkd2 is highly conserved in the clade of fission yeast species.

**Figure 8.**
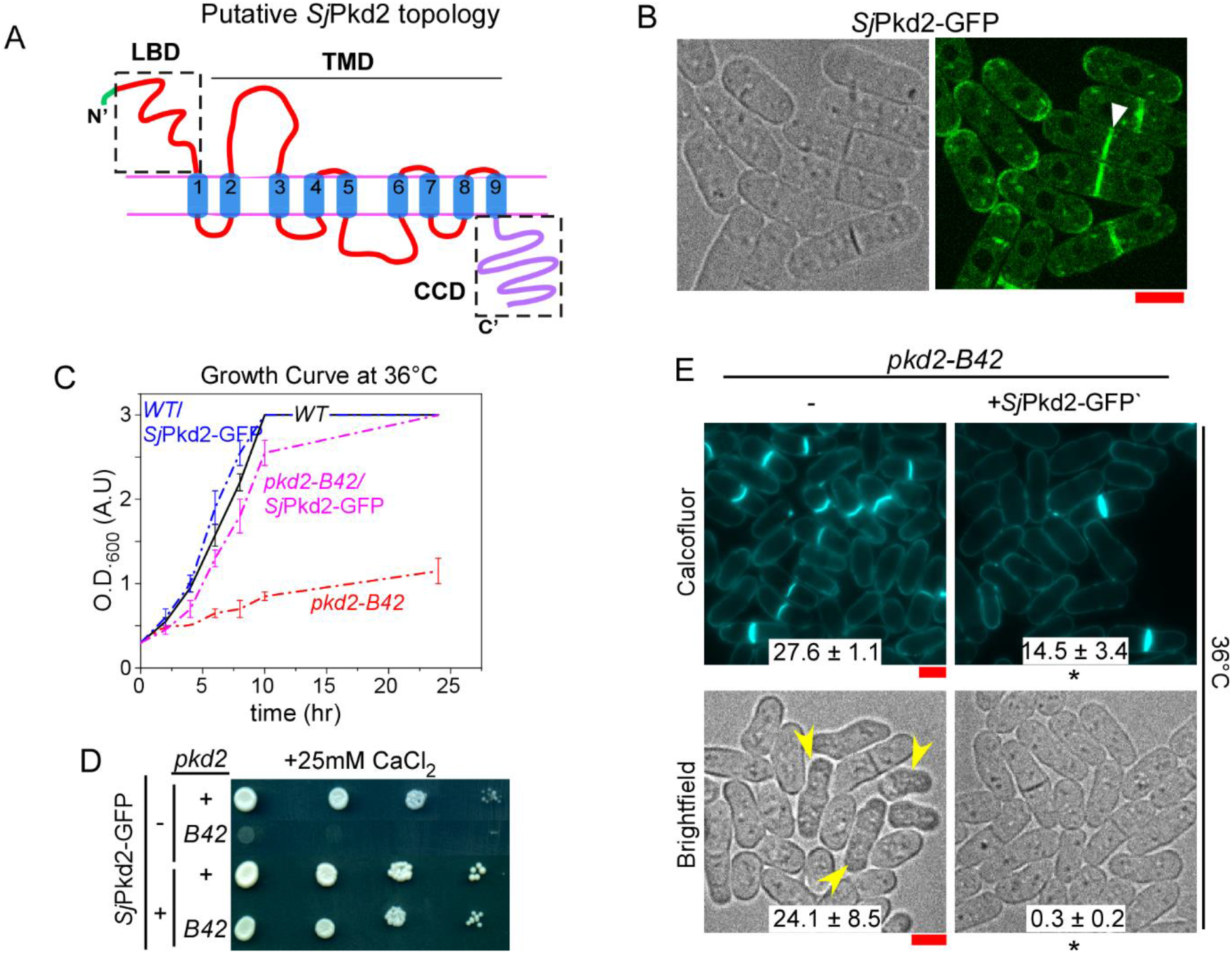
The function of Pkd2 is conserved in the *S. japonicus* homologue *Sj*Pkd2. **(A)** Diagram of the predicted topology of *Sj*Pkd2. Green: N-terminal signal peptide. **(B)** Micrographs of the wildtype *S. pombe* cells expressing *Sj*Pkd2-GFP (green) ectopically. Arrow: the equatorial division plane. **(C)** Growth curve of *S. pombe* cell cultures in YE5s liquid media. **(D)** Ten-fold dilution series of *S. pombe* cells grown on YE5S plus 25mM CaCl2 plates. **(E)** Micrographs of *pkd2-B42* mutant cells in the absence (left) or presence (right) of *Sj*Pkd2-GFP. Arrowhead: deflated cells identified in the bright field micrographs. The cells were inoculated at the restrictive temperature of 36°C for 4 hours. Number: the percentage of either septated (top) or deflated cells (bottom) (average ± standard deviations, n > 300). *: P ≤0.01 (Two-tailed student’s t-test). Data is pooled from at least two independent biological repeats. Scale: 5μm.

## Discussion

Despite what we have learned about the cellular functions and intracellular localization of Pkd2, little is known about its structure-function relationship. A structure of this putative ion channel would have greatly helped us to gain such insight, but it has not been experimentally determined. Solving the structure of such transmembrane proteins remains challenging, more so due to the likely oligomerization of Pkd2. Like most TRP channels, human polycystins assemble into a tetramer. With such constraint in mind, the Alphafold prediction, despite its imperfection, provided us with the best available option to dissect the function of each Pkd2 domain.

Of the three putative Pkd2 domains, we gained the most understanding on CCD through this study. Although it appeared to be mostly dispensable for the function of this essential protein, this cytoplasmic domain plays a critical role in regulating the localization of Pkd2. Without it, Pkd2 formed filament-like clusters on the plasma membrane. Such clustering is regulated by both eisosome and ER-PM contacts. In comparison to CCD, LBD and TMD are both essential for the function of Pkd2. They appear to be inseparable, likely owing to their extensive contacts with each other in the tertiary structure. Although similar structures can be found in two other fission yeast TRP channels, none of them can replace those of Pkd2. These two unique domains of Pkd2 are likely conserved in other fungi species.

Removal of CCD drastically altered the intracellular distribution of Pkd2 between the plasma membrane and the cytoplasm. With the presence of this mostly disordered domain, the full-length protein distributes extensively in the intracellular membrane structures, which likely include both endocytic vesicles and vacuoles, the yeast equivalent of lysosome. Such intracellular localization suggests that Pkd2 may be actively internalized through endocytosis followed by degradation in vacuoles. This process may require CCD as the initiation signal. This role of CCD could be achieved either through its direct ubiquitination or its interaction with an endocytic adaptor.

Without effective internalization, the fully functional Pkd2Δ CCD formed filament-like clusters on the plasma membrane, mostly along eisosomes. Clustering on the plasma membrane is common among the ion channels (Sato et al., 2019). Comparing the cell surface localization of Pkd2Δ CCD and Trp663, we may predict that Trp663 forms small clusters as well. However, toour knowledge, clustering of any other ion channel as short filaments in eisosomes, as Pkd2Δ CCD does, has not been reported before. Eisosomes/MCCs (membrane compartment of Can1) are furrowing microdomains of the yeast plasma membrane (Malinska et al., 2003; Young et al., 2002). Although Pkd2 may be the only ion channel in fission yeast eisosomes, many amino acid-polyamine-organocation (APC) transporters have been found there (Bianchi et al., 2018; Gournas et al., 2018; Moharir et al., 2018).

Just like the exact function of eisosomes remains debated, the importance of Pkd2 clustering in eisosome is still unclear. Two likely roles of eisosome have emerged (Appadurai et al., 2020). First, they could prevent those APC transporters from being endocytosed and degraded (Grossmann et al., 2008). Secondly, this raft-like microdomain could provide the membrane reservoir when the cells need to expand their plasma membrane under mechanical stress (Kabeche et al., 2015; Lemière et al., 2021). Both could potentially explain the clustering of Pkd2 in eisosomes. The first is consistent with the reduced internalization of Pkd2 in the *pkd2Δ CCD* mutant. As for the second, the storage in eisosomes may serve as a mechanism to prevent Pkd2 from being activated as an ion channel outside of the growth zones.

Surprisingly, disruption of ER-PM contacts, another microdomain of the plasma membrane, exacerbated clustering of Pkd2. The ER-PM contacts are well-established as sites of lipid transfer between ER and the plasma membrane (Manford et al., 2012). One likely explanation for this observation is that Pkd2Δ CCD may form small clusters at the ER-PM contact sites. Thus, reduction of ER-PM contacts in *scs2Δ scs22Δ* mutant may have made more Pkd2 available to form filament-like clusters in eisosomes.

The predicted structure of LBD is unique in its Ig-like fold with a potential for lipid-binding. To our knowledge, the arrangement of its eight β-strands is unlike any other Ig-like domain found so far (Bork et al., 1994). Sequence analysis had suggested that LBD may be a ML-like (MD-2-related lipid-recognition) domain found in Der f 2, a major mite allergen (Ichikawa et al., 1998) or NPC2 protein (Niemann-Pick disease type C2) (Friedland et al., 2003). Both are capable of binding lipids (Johannessen et al., 2005). However, such a prediction is contradictory to the projected extracellular position of LBD. Alternatively, LBD may gate the ion channel pore of Pkd2 channel, like the conserved TOP domain of human polycystins (Shen et al., 2016). This explanation would agree with the Alphafold prediction which shows LBD sits on the top of TMD and makes extensive contacts.

In summary, we combined the predicted tertiary structure of the essential fission yeast channel Pkd2 with genetic analyses to dissect its domains for the first time. The putative structure is likely conserved in both fission yeast and other fungi in which Pkd2 is a highly conserved essential protein (Hsiang and Baillie, 2005). Our novel discovery of the clustering of this putative ion channel in eisosomes indicates that its distribution in the plasma membrane is tightly regulated through the cytoplasmic tail, by the endocytosis-mediated internalization and the organization of the plasma membrane.

## Materials and methods

### Structure prediction

We used ChimeraX (UCSF) to retrieve the predicted tertiary structures from Alphafold. The Uniprot numbers for Pkd2, Trp663, Trp1322 and *Sj*Pkd2 are Q09917, O74520, O94543 and B6K7F8 respectively. All the structures in the figures are also rendered using ChimeraX.

### Yeast genetics and cell culture

We followed the standard protocols for yeast cell culture and genetics (Moreno et al., 1991). Fission yeast tetrads were sporulated on SPA5S plates and dissected using a SporePlay+ dissection microscope (Singer, UK). Transformation of yeast used the standard lithium-acetate method. All the strains used in this study are listed in Supplemental Table S3.

For assays of viability, overnight cultures were diluted and grown for additional six hours before being spotting for the ten-fold dilution series. To measure growth curve in liquid culture, we diluted overnight cultures to optical density of 0.3 at A_600nm_. We then measured the cell density with a spectrometer (Eppendorf) every 2 hours for the next 10 hours followed by the final measurement at 24 hours.

### Cloning of Pkd2 truncation mutants and chimeras

For molecular cloning, both the gene of interest and the vector were amplified by PCR using Q5 polymerase (New England Biolabs, US). After being digested with DpnI, the PCR products were purified using NucleoSpin^®^ Gel and PCR Clean-up kit (Macherey-Nagel #740609). Purified DNA was cloned into pAV0714 (Vještica et al., 2020) using NEBuilder^®^ HiFi DNA assembly kit (NEB #E2621). After being verified by Sanger sequencing, 2μg plasmid was linearized with the restriction enzyme AfeI. The constructs tagged with mCherry are integrated at *ade6* locus and linearized by PmeI. The linearized vectors were purified and then transformed into yeast using lithium-acetate method. mEGFP was used to tag the Pkd2 truncation mutants at their endogenous locus or from *leu1* locus. sfGFP (super-fold) version was used for those expressed form the *ura4* locus.

Most over-expression constructs used in this study were first cloned into pAV0714 (Vještica et al., 2020) driven by a constitutively active actin promoter *Pact*. at *ura4* locus. They were integrated into the yeast genome. The lone exception is LBD-GFP which is expressed at *leu1* locus driven by *3nmt1* promoter. To construct LBD-GFP expression vector, the DNA sequence of LBD including that of the signal peptide was amplified and cloned into the vector pFA-kanMX6-3nmt1GFP using HiFi assembly kit. The DNA fragment encoding P3nmt1-LBD-GFP was then integrated into *leu1* locus through PCR-based homologous recombination (Bähler et al., 1998).

### Microscopy

For microscopy, 1-2 ml of exponentially growing yeast cells at 25°C with a density between 5.0 × 10^6^/ml and 1.0 × 10^7^/ml in YE5s liquid media (unless specified), were harvested by centrifugation at 4000 rpm for 1 min and resuspended in 50 μl YE5s. Resuspended cells (6 μl) were spotted to a 25% gelatin + YE5s pad and sealed under the coverslip with VALAP (1:1:1 of Vaseline, Lanolin, and Paraffin) (Wang et al., 2016).

Live cell microscopy was carried out on an Olympus IX71 microscope. It is equipped with a 100x (NA = 1.40) and 60x (NA = 1.40) objective lenses, a confocal spinning-disk unit (CSU-X1; Yokogawa, Japan), a motorized XY stage with a Piezo Z Top plate (ASI), in a designated microscopy room where the temperature was maintained at around 22 ± 2°C. The images were captured on an Ixon-897 EMCCD camera controlled by iQ3.0 (Andor, Ireland). Solid-state lasers of 488 and 561 nm were used for imaging green and red fluorescence respectively. For time-lapse microscopy, the cells were imaged for two hours with 2-minutes interval and with Z-series of 15 slices at a step size of 0.5 μm. To help minimize the potential variations in culture and microscopy conditions, the wild-type control and the experimental groups were imaged on the same day. 60x objective was used to take brightfield images for the quantification of deflated cells.

For calcofluor staining, 2 ml of exponentially growing cells were harvested at 4000rpm for 1 min. The cells were fixed in 4% paraformaldehyde while being rotated for 10 mins. We spun down the fixed cells and washed them three times with 1 ml of TEMK buffer. The washed cells were re-suspended in 1ml TEMK buffer and stored at 4ºC until next day. Just before imaging, 1 μl of calcofluor (1μg/ml) (Sigma #18909) was added to the 1 ml of cells. The cells were stained while being rotated on a rocker in the dark for 5 minutes before they were pelleted and resuspend in 50 μl TEMK. The stained sample (5 μl) was spotted directly on a glass slide, covered with cover slip, and sealed with VALAP. Images were taken at 60x using an epifluorescence microscope Olympus IX81 equipped with a mercury lamp and a digital camera.

### Image processing and analysis

NIH ImageJ was used for processing all the images with either freely available or customized macros/plug-ins. For quantitative analysis of the fluorescence micrographs of a time-lapse series, we first corrected X-Y drifting using StackReg (Thevenaz et al., 1998). Average intensity projections of Z-slices were used for quantification. Kymographs of the time-series of the contractile rings were generated to determine both the start and end of ring closure.

To quantify the colocalization between two proteins, maximum intensity projections of the two top slices of a full Z-series where the green fluorescence was visible were generated. ImageJ (Fiji) macro Coloc2 was used to quantify the Pearson’s correlation coefficient (R) (no threshold). R value <0.1 is considered as no colocalization, 0.1-0.5 as partial and >0.5 is considered as significant colocalization.

## Abbreviations

(LBD): Lipid Binding Domain
(TMD): Transmembrane domain
(CCD): Cytoplasmic coiled-coil domain
(ER): Endoplasmic reticulum
(PM): Plasma membrane
(TRP): Transient Receptor Potential
(VAPs): Vesicle-associated membrane protein-associated proteins
(YE5S): Yeast extract with 5 supplements
(SPA5S): Sporulation agar with 5 supplements
(TEMK): Tris-HCl EGTA Magnesium Potassium
(a.a): Amino acid
(WT): Wild-type

## Author contributions

QC conceptualized the study; QC, MM and DS designed the experiments; QC, MM, DS, BB and PC acquired and analyzed the data; QC and MM wrote the manuscript.

## Acknowledgements

We would like to thank Dr. Zhang Dan (Temasek Life Sciences Laboratory, Singapore), Dr. Snezhana Oliferenko (King’s College, London) and Dr. James Moseley (Geisel School of Medicine at Dartmouth, New Hampshire) for sharing their yeast strains, and Dr. Sophie Martin (University of Lausanne, Switzerland) for sharing the pAV0714 plasmid. We thank Dr. Steven Chou (University of Connecticut) for assisting with our structure prediction using Alphafold. We acknowledge National BioResource Project (NBRP)-Yeast (Japan) for sending the yeast strains. We also thank the Chen lab members Abhishek Poddar and Clare Muller at The University of Toledo for technical assistance. This work has been supported by the National Institutes of Health grant R15GM134496 and the Nation Science Foundation grant 2144701 to QC. The content is solely the responsibility of the authors and does not necessarily represent the official views of the National Institutes of Health. The authors declare no conflicts of interests.

## Supplemental Materials

### Supplemental Tables

**Supplemental Table S1.** Pkd2 truncation constructs used in this study.

| Name | Amino acid sequence | Locus | Intracellular localization | Rescue of *pkd2-B42* |
| --- | --- | --- | --- | --- |
| Pkd2-GFP | 1-710 | *ura4* | PM + intracellular membrane structures | Yes |
| Pkd2(LBD)-GFP | 1-170 | *leu1* | Cytoplasmic | N/A |
| Pkd2(TMD)-GFP | (1-23) + (171-575) | *ura4* | Cortical ER | No |
| Pkd2(1-TM1)-mCherry | 1-193 | *ade6* | Cortical ER | No |
| Pkd2(1-TM3)-GFP | 1-360 | *ura4* | Cortical ER | No |
| Pkd2(1-TM5)-GFP | 1-430 | *ura4* | Cortical ER | No |
| Pkd2(1-TM7)-GFP | 1-515 | *ura4* | Cortical ER | No |
| Pkd2(1-TM8)-GFP | 1-550 | *ura4* | Cortical ER | No |
| Pkd2(1-TM1-TM4-9)-GFP | (1-170) + (205-583) | *ura4* | Cortical ER | No |
| Pkd2(1-TM1-TM8-9)-GFP | (1-170) + (523-583) | *ura4* | Cortical ER | No |
| Pkd2(CCD)-GFP | (576-710) | *ura4* | Cytoplasmic | N/A |

PM: plasma membrane; TM: transmembrane helix; N/A: not applicable

**Supplemental Table S2.** Septation and deflation indices of the cells expressing Chimera I.

| Strain | Ectopically expressed construct | Septated cells (%) | Deflated cells (%) |
| --- | --- | --- | --- |
| *WT* | - | 13.3±1.3 | 0.1±0.1 |
|  | Pkd2 | 13.8±2.4 | 0.2±0.2 |
|  | Chimera I | 12.2±3.8 | 0.1±0.1 |
| *pkd2-B42* | - | 27.6±1.1 | 24.1±8.5 |
|  | Pkd2 | 10.1±2.5 | 0.3±0.05 |
|  | Chimera I | 16±3.3 | 1.0±0.3 |

-: no construct
Data is pooled from at least two independent biological repeats.

**Supplemental Table S3.** Yeast strains used in this study.

| Strains | Genotype | Source |
| --- | --- | --- |
| FY527 | *h- leu1-32 ura4-D18 his3-D1 ade6-M216* | Lab stock |
| QC-Y690 | *h- pkd2-GFP-kanMX6 leu1-32 ura4-D18 his3-D1 ade6-M216* | Lab stock |
| QC-Y693 | *h- pkd2-GFP-kanMX6 rlc1-tdTomato-NatMX6* | Lab stock |
| QC-Y758 | *h- pkd2::pkd2(ΔCCD)-GFP-KanMX6 leu1-32 ura4-D18 his3-D1 ade6-M216* | Lab stock |
| QC-Y771 | *h? pkd2::pkd2(ΔCCD)-GFP-KanMX6 rlc1-tdTomato-NatMX6* | Lab stock |
| QC-Y988 | *h+ trp663::kanMX6* | Lab stock |
| QC-Y1031 | *h- pkd2::pkd2-B42-ura4+his5+ leu1-32* | Lab stock |
| QC-Y1032 | *h+ pkd2::pkd2-B42-ura4+his5+ leu1-32* | Lab stock |
| QC-Y1058 | *h- leu1::P3nmt1-Pkd2(LBD)-GFP-KanMX6 ura4-D18 his3-D1 ade6-M216* | This study |
| QC-Y1117 | *h- ura4::Pact1-Pkd2-GFP-ura4+ leu1-32 his3-D1 ade6-M216* | This study |
| QC-Y1119 | *h? pkd2::pkd2-B42 ura4::Pact1-Pkd2-GFP-ura4+ his5+ leu1-32 his3-D1 ade6-M216* | This study |
| QC-Y1167 | *h- ura4::Pact1-Pkd2<sup>LBD</sup>-PC1<sup>TMD</sup>-GFP-ura4+ leu1-32 his3-D1 ade6-M216* | This study |
| QC-Y1184 | *h? pkd2::pkd2-B42 ura4::Pact1- Pkd2<sup>LBD</sup>-PC1<sup>TMD</sup> -GFP-ura4+ his5+ leu1-32 his3-D1 ade6-M21X* | This study |
| QC-Y1297 | *h+ade6::Pact1-Pkd2(1-TM1)-mCherry-ade6+ leu1-32 ura4-D18 his3-D1* | This study |
| QC-Y1298 | *h+ ade6::Pact1-Pkd2(LBD+TM1)-PC2<sup>TMD</sup> -mCherry-ade6+ leu1-32 ura4-D18 his3-D1* | This study |
| QC-Y1303 | *h? ura4::Pact1- Pkd2<sup>LBD</sup>-PC1<sup>TMD</sup> -GFP-ura4+ade6::Pact1- Pkd2(LBD+TM1)-PC2<sup>TMD</sup> -mCherry-ade6+ leu1-32 ura4-D18 his3-D1* | This study |
| QC-Y1315 | *h? pkd2::pkd2-B42 ade6::Pact1- Pkd2(LBD+TM1)-PC2<sup>TMD</sup> -mCherry-ade6+-ura4+his5+ leu1-32* | This study |
| QC-Y1318 | *h? pkd2::pkd2-B42 ura4::Pact1- Pkd2<sup>LBD</sup>-PC1<sup>TMD</sup> -GFP ade6::Pact1- Pkd2(LBD+TM1)-PC2<sup>TMD</sup> -mCherry-ade6+ ura4+ his5+ leu1-32* | This study |
| QC-Y1336 | *h- ura4::Pact1-Pkd2(1-TM3)-GFP-ura4+ leu1-32 his3-D1 ade6-M216* | This study |
| QC-Y1337 | *h- ura4::Pact1-Pkd2(1-TM5)-GFP-ura4+ leu1-32 his3-D1 ade6-M216* | This study |
| QC-Y1338 | *h- ura4::Pact1-Pkd2(1-TM7)-GFP-ura4+ leu1-32 his3-D1 ade6-M216* | This study |
| QC-Y1339 | *h+ trp1322::KanMX6* | Lab stock |
| QC-Y1344 | *h? pkd2::pkd2-B42 ura4::Pact1-Pkd2(1-TM3)-GFP -ura4+ his5+ leu1-32 his3-D1 ade6-M21X* | This study |
| QC-Y1345 | *h? pkd2::pkd2-B42 ura4::Pact1-Pkd2(1-TM5)-GFP-ura4+ -ura4+ his5+ leu1-32 his3-D1 ade6-M21X* | This study |
| QC-Y1346 | *h? pkd2::pkd2-B42 ura4::Pact1-Pkd2(1TM7)-GFP-ura4+ his5+ leu1-32 his3-D1 ade6-M21X* | This study |
| QC-Y1360 | *h- ura4::Pact1-Pkd2(1-TM8)-GFP-ura4+ leu1-32 his3-D1 ade6-M216* | This study |
| QC-Y1364 | *h? trp1322::KanMX6 pkd2::pkd2-B42-ura4+ his5+ leu1-32* | Lab stock |
| QC-Y1367 | *h?pkd2::pkd2-B42 ura4::Pact1-Pkd2(1-TM8)-GFP-ura4+ his5+ leu1-32 his3-D1 ade6-M21X* | This study |
| QC-Y1375 | *h- ura4::Pact1-Trp663<sup>LBD</sup> Pkd2<sup>(TMD+CCD)</sup>-GFP-ura4+ leu1-32 his3-D1 ade6-M216* |  |
| QC-Y1376 | *h- ura4::Pact1-Pkd2(1-TM1-TM4-9)-GFP-ura4+ leu1-32 his3-D1 ade6-M216* | This study |
| QC-Y1377 | *h- ura4::Pact1-Pkd2(1-TM1-TM8-9)-GFP-ura4+ leu1-32 his3-D1 ade6-M216* | This study |
| QC-Y1386 | *h? pkd2::pkd2-B42 ura4::Pact1-Pkd2(1-TM1-TM4-9)-GFP-ura4+ +his5+ leu1-32 his3-D1 ade6-M21X* | This study |
| QC-Y1387 | *h? pkd2::pkd2-B42 ura4::Pact1-Pkd2(1-TM1-TM8-9)-GFP-ura4+ his5+ leu1-32 his3-D1 ade6-M21X* | This study |
| QC-Y1391 | *h? pkd2::pkd2-B42 ura4::Pact1- Trp663<sup>LBD</sup> Pkd2<sup>(TMD+CCD)</sup>-GFPura4+his5+ leu1-32 his3-D1 ade6-M21X* |  |
| QC-Y1404 | *h? trp663::kanMX6 pkd2::pkd2-B42-ura4+his5+ leu1-32* | This study |
| QC-Y1405 | *h- ura4::Pact1-Trp663-GFP-ura4+ leu1-32 his3-D1 ade6-M216* | This study |
| QC-Y1411 | *h? pkd2::pkd2-B42 ura4::Pact1-Trp663-GFP-ura4+his5+ leu1-32 his3-D1 ade6-M21X* | This study |
| QC-Y1419 | *h- ura4::Pact1-Trp1322-GFP-ura4+ leu1-32 his3-D1 ade6-M216* | This study |
| QC-Y1428 | *h? pkd2::pkd2-B42 ura4::Pact1-Trp1322-GFP -ura4+ his5+ leu1-32 his3-D1 ade6-M21X* | This study |
| QC-Y1438 | *h- ura4::Pact1- Pkd2<sup>LBD</sup> Trp663<sup>(TMD+CCD)</sup>-GFP-ura4+ leu1-32 his3-D1 ade6-M216* | This study |
| QC-Y1442 | *h? pkd2::pkd2-B42-ura4 ura4::Pact1-Pkd2<sup>LBD</sup>Trp663<sup>(TMD-CCD)</sup>-GFP + his5+ leu1-32 ade6-M21X* | This study |
| QC-Y1443 | *h? ura4::Pact1- Trp663<sup>LBD</sup> Pkd2<sup>(TMD+CCD)</sup>-GFP-ura4+ leu1::Pbip1-mCherry-ADEL ade6? leu1?* | This study |
| QC-Y1444 | *h? ura4::Pact1- Pkd2<sup>LBD</sup> Trp663<sup>(TMD+CCD)</sup>-GFP-ura4+ leu1::Pbip1-mCherry-ADEL ade6? leu1?* | This study |
| QC-Y1460 | *h- ura4::Pact1-SjPkd2-GFP-ura4+ leu1-32 his3-D1 ade6-M216* | This study |
| QC-Y1467 | *h? ura4::Pact1-Pkd2<sup>(TMD)</sup>-GFP-ura4+ leu1::Pbip1-mCherry-ADEL ade6X leu1X* | This study |
| QC-Y1473 | *h? pkd2::pkd2-B42 ura4::Pact1-SjPkd2-GFP -ura4+ his5+ leu1-32 ade6-M21X* | This study |
| QC-Y1498 | *h+ pkd2::pkd2(ΔCCD)-GFP-KanMX6 scs2::ura4+ scs22::ura4+ ura4-D18 leu1-32 his3-D1 ade6?* | This study |
| QC-Y1518 | *h? pkd2-GFP-kanMX6 scs2::ura4+ scs22::ura4+ ura4-D18 leu1-32 his3-D1 ade6?* | This study |
| QC-Y1521 | *h? pkd2::pkd2(ΔCCD)-GFP-KanMX6 pil1-mCherry::natR* | This study |
| QC-Y1527 | *h+/h pkd2::KanMX6/+ ura4::Pact1-Trp663<sup>LBD</sup> Pkd2<sup>(TMD+CCD)</sup>-GFP-ura4+* | This study |
| QC-Y1530 | *h? pkd2::pkd2(ΔCCD)-GFP-KanMX6 scs2::ura4+ scs22::ura4+ pil1-mCherry::natR* | This study |
| QC-Y1534 | *h- ura4::Pact1 Pkd2(CCD)-GFP-ura4+leu1-32 his3-D1 ade6-M216* | This study |
| QC-Y1537 | *h? pkd2::KanMX6 ura4::Pact1-Pkd2-GFP-ura4+* | This study |
| QC-Y1545 | *h? pkd2::pkd2-GFP-KanMX6 pil1-mCherry::natR* | This study |

### Supplemental Figures

**Supplemental Figure S1.**
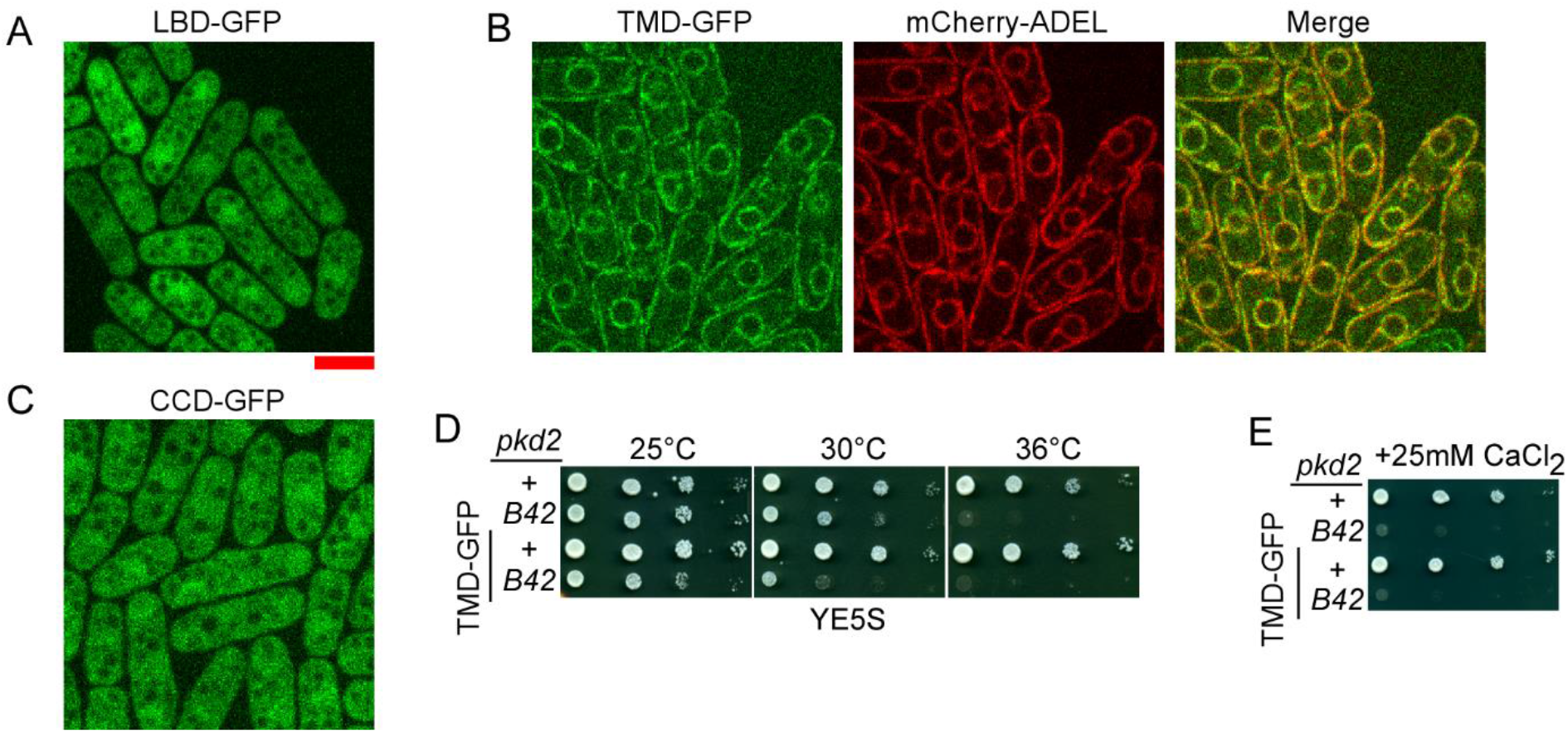
Localization and function of individual Pkd2 domains. **(A-C)** Micrographs of the cells ectopically expressing either LBD-GFP (A), or TMD-GFP (B), or CCD-GFP (C). mCherry-ADEL was co-expressed as the marker for the cortical ER. The center slice of the Z-series is shown. Scale: 5μm. **(D-E)** Ten-fold dilution series of yeast expressing TMD-GFP on either YE5S (D) or YE5S plus 25mM CaCl_2_ plates (E).

**Supplemental Figure S2.**
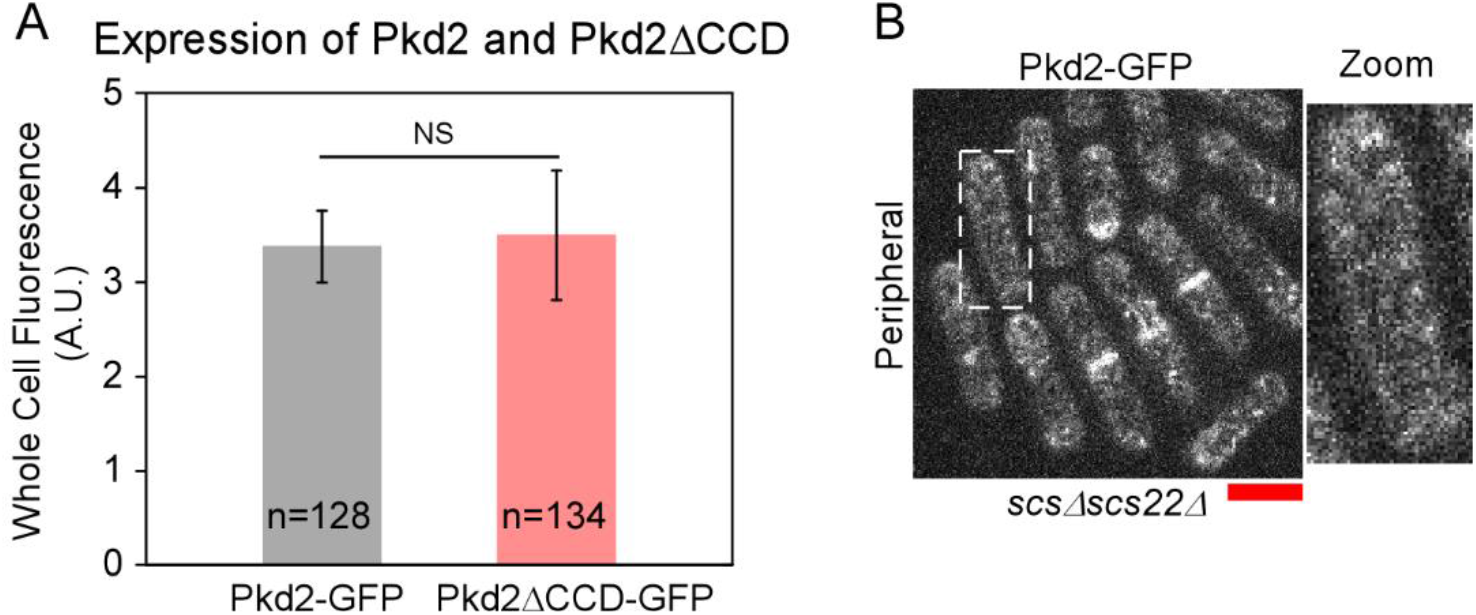
Expression and localization of Pkd2 and Pkd2ΔCCD. **(A)** Bar graph of the average whole-cell fluorescence of the cells expressing either Pkd2-GFP or Pkd2Δ CCD-GFP from the endogenous locus. NS: no significant differences (Two-tailed student’s t-test). **(B)** Micrograph of *scs2Δscs22Δ* cells expressing Pkd2-GFP. Right: Magnified view of a representative cell (dashed line). Scale: 5μm.

**Supplemental Figure S3.**
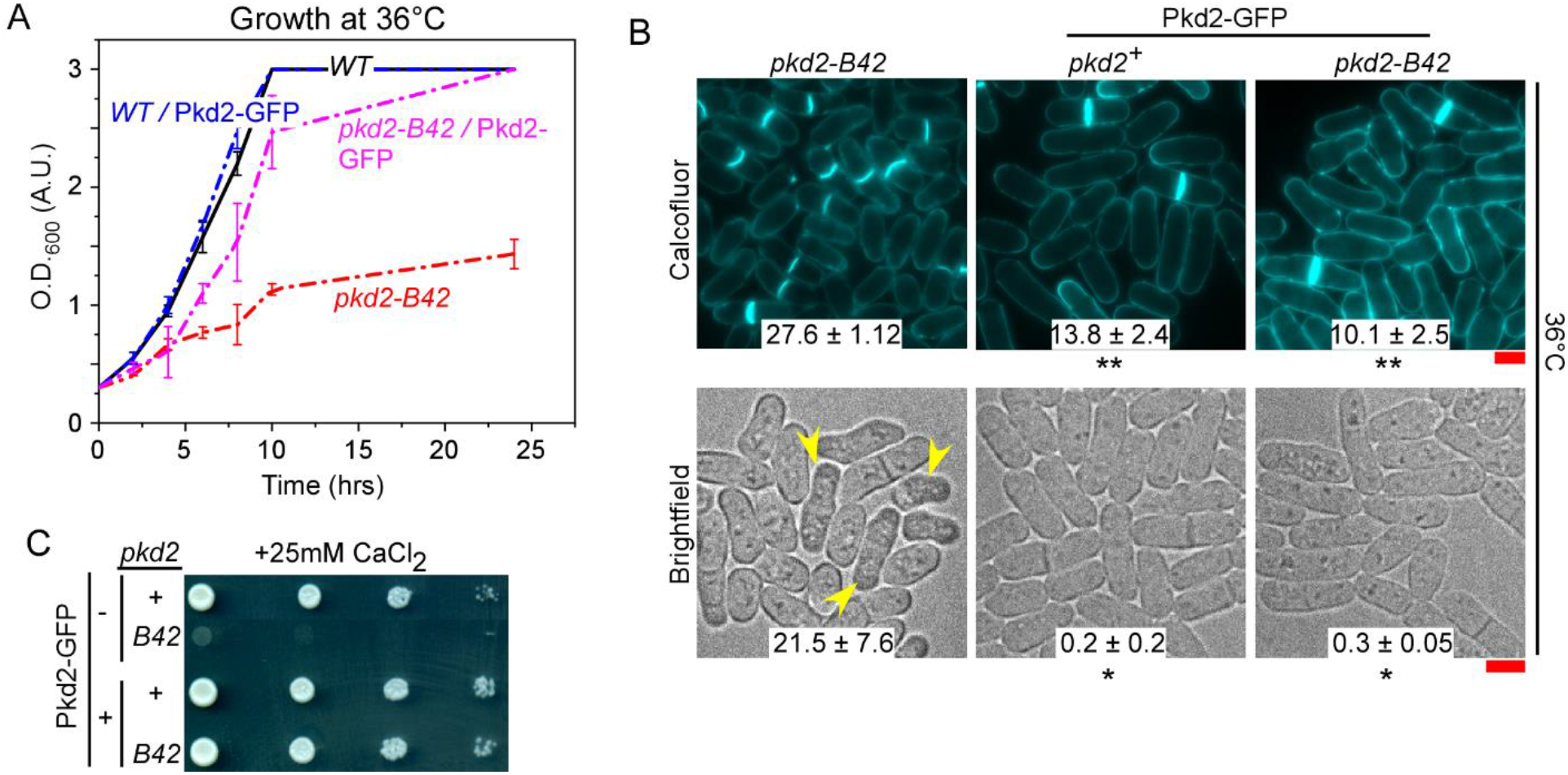
Ectopic expression of full-length Pkd2 is fully functional. **(A)** Growth curve of yeast in liquid culture at the restrictive temperature 36°C. **(B)** Micrographs of *pkd2-B42*, and wild-type or *pkd2-B42* cells in presence of ectopic Pkd2-GFP cells either fixed and stained with calcofluor (top) or imaged live (bottom). The cells were inoculated at 36°C for 4 hours before fixation or being imaged live. Number: the average percentage of septated (top) or deflated cells (bottom) (n > 300). *pkd2-B42* data are from Fig. 8E. **(C)** Ten-fold dilution series of yeast on YE5S plus 25mM CaCl_2_ plates. *: P ≤0.01, **: P≤0.001 (Two-tailed student’s t-test). Data is from at least two independent biological repeats. Scale: 5μm

**Supplemental Figure S4.**
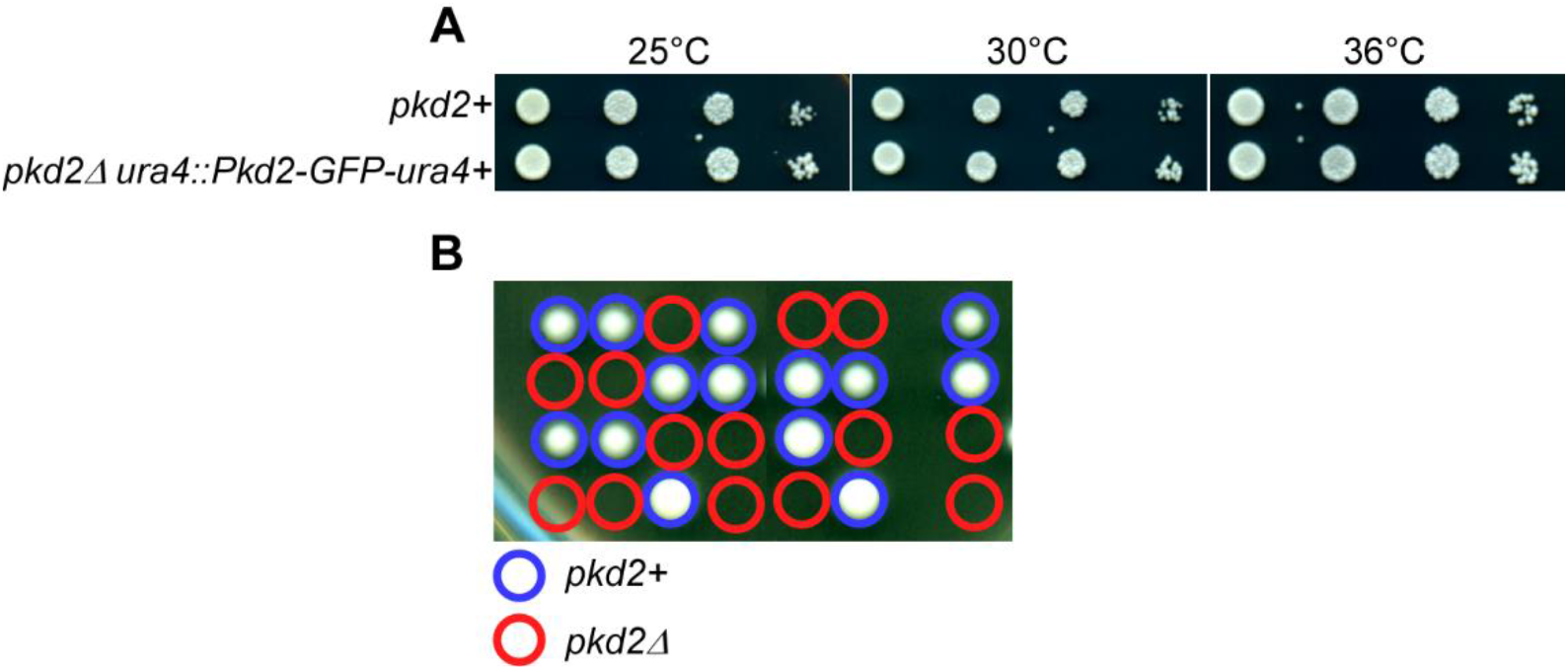
Ectopic expression of full-length Pkd2 replaces the endogenous gene, but the chimera between Pkd2 and Trp663 can’t. **(A)** Ten-fold dilution series of yeast on YE5S. **(B)** Tetrad-dissection plate of sporulated *pkd2+/pkd2Δ ura4::Trp663*^*LBD*^*Pkd2*^*(TMD-CCD)*^*-GFP-ura4+*. The viable cells have *pkd2+* (blue circles).The inviable cells have *pkd2Δ* (red circles).

**Supplemental Figure S5.**
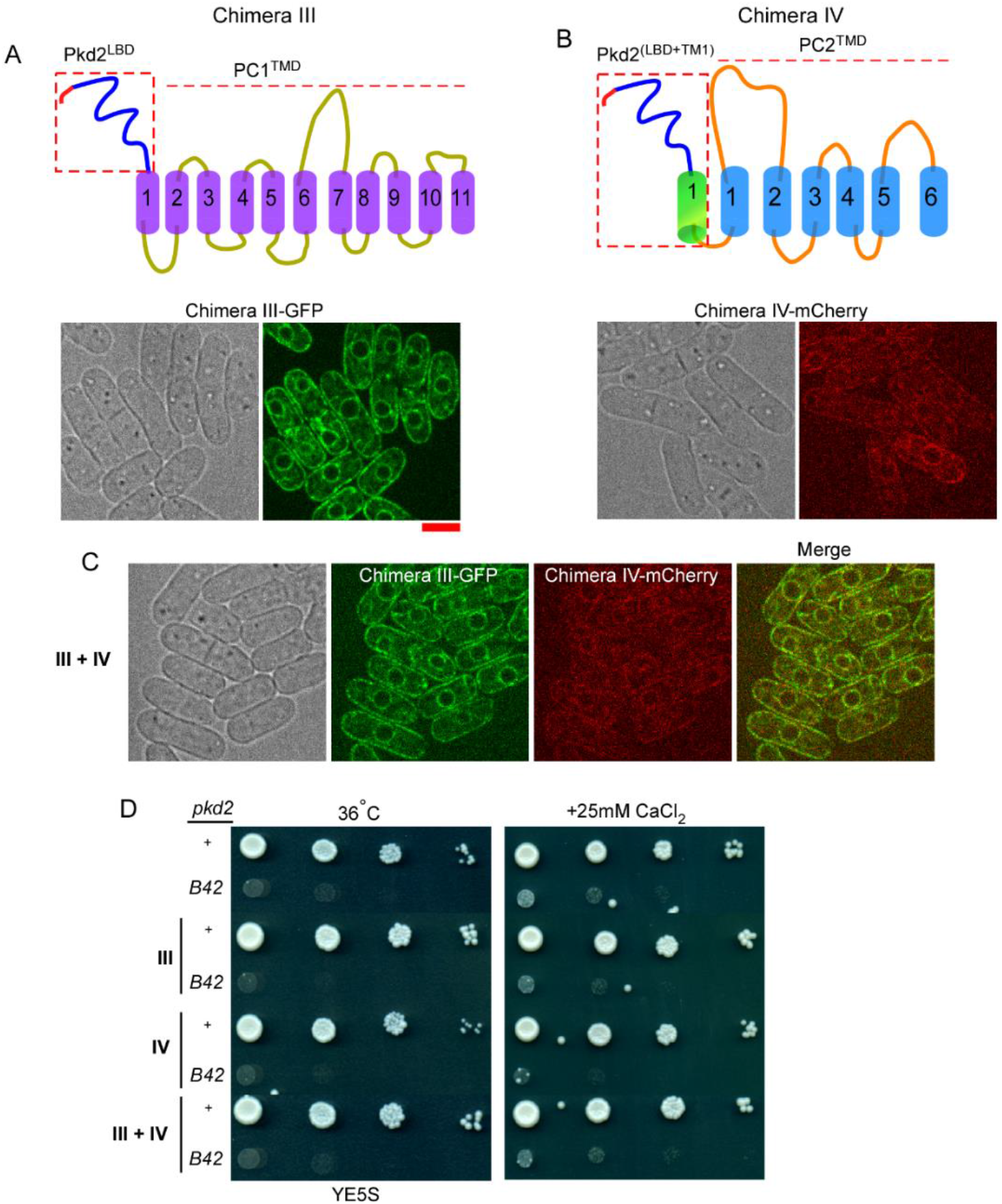
TMD of Pkd2 is not interchangeable with those of either human polycystins PC1 or PC2. **(A-B)** Putative topology of Chimera III, Pkd2^LBD^-PC1^TMD^ (A) and Chimera IV, Pkd2^(LBD+TM1)^ -PC2^TMD^ (B). **(C)** Micrographs of the cells expressing both Chimera III tagged with GFP (green), and Chimera IV tagged with mCherry (red). **(D)** Ten-fold dilution series of yeast on YE5S plates at 36°C and YE5S plus 25mM CaCl_2_ plates. Scale: 5μm.

## References

Appadurai, D., L. Gay, A. Moharir, M.J. Lang, M.C. Duncan, O. Schmidt, D. Teis, T.N. Vu, M. Silva, E.M. Jorgensen, and M. Babst. 2020. Plasma membrane tension regulates eisosome structure and function. Molecular biology of the cell. 31:287–303.

Bähler, J., J.Q. Wu, M.S. Longtine, N.G. Shah, A. Mckenzie III, A.B. Steever, A. Wach, P. Philippsen, and J.R. Pringle. 1998. Heterologous modules for efficient and versatile PCR-based gene targeting in Schizosaccharomyces pombe. Yeast. 14:943–951.

Bianchi, F., L. Syga, G. Moiset, D. Spakman, P.E. Schavemaker, C.M. Punter, A.B. Seinen, A.M. van Oijen, A. Robinson, and B. Poolman. 2018. Steric exclusion and protein conformation determine the localization of plasma membrane transporters. Nature communications. 9:501.

Bork, P., L. Holm, and C. Sander. 1994. The immunoglobulin fold. Structural classification, sequence patterns and common core. Journal of molecular biology. 242:309–320.

Foggensteiner, L., A.P. Bevan, R. Thomas, N. Coleman, C. Boulter, J. Bradley, O. Ibraghimov-Beskrovnaya, K. Klinger, and R. Sandford. 2000. Cellular and subcellular distribution of polycystin-2, the protein product of the PKD2 gene. Journal of the American Society of Nephrology. 11:814–827.

Friedland, N., H.-L. Liou, P. Lobel, and A.M. Stock. 2003. Structure of a cholesterol-binding protein deficient in Niemann–Pick type C2 disease. Proceedings of the National Academy of Sciences. 100:2512–2517.

Geng, L., Y. Segal, B. Peissel, N. Deng, Y. Pei, F. Carone, H.G. Rennke, A.M. Glücksmann-Kuis, M.C. Schneider, and M. Ericsson. 1996. Identification and localization of polycystin, the PKD1 gene product. The Journal of clinical investigation. 98:2674–2682.

Gournas, C., S. Gkionis, M. Carquin, L. Twyffels, D. Tyteca, and B. Andre. 2018. Conformation-dependent partitioning of yeast nutrient transporters into starvation-protective membrane domains. Proceedings of the National Academy of Sciences of the United States of America. 115:E3145–E3154.

Grieben, M., A.C. Pike, C.A. Shintre, E. Venturi, S. El-Ajouz, A. Tessitore, L. Shrestha, S. Mukhopadhyay, P. Mahajan, and R. Chalk. 2017. Structure of the polycystic kidney disease TRP channel Polycystin-2 (PC2). Nature structural & molecular biology. 24:114–122.

Grossmann, G., J. Malinsky, W. Stahlschmidt, M. Loibl, I. Weig-Meckl, W.B. Frommer, M. Opekarova, and W. Tanner. 2008. Plasma membrane microdomains regulate turnover of transport proteins in yeast. The Journal of cell biology. 183:1075–1088.

Hsiang, T., and D.L. Baillie. 2005. Comparison of the yeast proteome to other fungal genomes to find core fungal genes. J Mol Evol. 60:475–483.

Ichikawa, S., H. Hatanaka, T. Yuuki, N. Iwamoto, S. Kojima, C. Nishiyama, K. Ogura, Y. Okumura, and F. Inagaki. 1998. Solution structure of Der f 2, the major mite allergen for atopic diseases. The Journal of biological chemistry. 273:356–360.

Johannessen, B.R., L.K. Skov, J.S. Kastrup, O. Kristensen, C. Bolwig, J.N. Larsen, M. Spangfort, K. Lund, and M. Gajhede. 2005. Structure of the house dust mite allergen Der f 2: implications for function and molecular basis of IgE cross-reactivity. FEBS letters. 579:1208–1212.

Jumper, J., R. Evans, A. Pritzel, T. Green, M. Figurnov, O. Ronneberger, K. Tunyasuvunakool, R. Bates, A. Žídek, and A. Potapenko. 2021. Highly accurate protein structure prediction with AlphaFold. Nature. 596:583–589.

Kabeche, R., S. Baldissard, J. Hammond, L. Howard, and J.B. Moseley. 2011. The filament-forming protein Pil1 assembles linear eisosomes in fission yeast. Molecular biology of the cell. 22:4059–4067.

Kabeche, R., L. Howard, and J.B. Moseley. 2015. Eisosomes provide membrane reservoirs for rapid expansion of the yeast plasma membrane. Journal of cell science. 128:4057–4062.

Lamas, I., L. Merlini, A. Vjestica, V. Vincenzetti, and S.G. Martin. 2020. Optogenetics reveals Cdc42 local activation by scaffold-mediated positive feedback and Ras GTPase. PLoS biology. 18:e3000600.

Lemière, J., Y. Ren, and J. Berro. 2021. Rapid adaptation of endocytosis, exocytosis, and eisosomes after an acute increase in membrane tension in yeast cells. Elife. 10:e62084.

Liu, X., T. Vien, J. Duan, S.-H. Sheu, P.G. DeCaen, and D.E. Clapham. 2018. Polycystin-2 is an essential ion channel subunit in the primary cilium of the renal collecting duct epithelium. Elife. 7:e33183.

Malinska, K., J. Malinsky, M. Opekarova, and W. Tanner. 2003. Visualization of protein compartmentation within the plasma membrane of living yeast cells. Molecular biology of the cell. 14:4427–4436.

Manford, A.G., C.J. Stefan, H.L. Yuan, J.A. Macgurn, and S.D. Emr. 2012. ER-to-plasma membrane tethering proteins regulate cell signaling and ER morphology. Developmental cell. 23:1129–1140.

Moharir, A., L. Gay, D. Appadurai, J. Keener, and M. Babst. 2018. Eisosomes are metabolically regulated storage compartments for APC-type nutrient transporters. Molecular biology of the cell. 29:2113–2127.

Moreno, S., A. Klar, and P. Nurse. 1991. Molecular genetic analysis of fission yeast Schizosaccharomyces pombe. Methods in enzymology. 194:795–823.

Morris, Z., D. Sinha, A. Poddar, B. Morris, and Q. Chen. 2019. Fission yeast TRP channel Pkd2p localizes to the cleavage furrow and regulates cell separation during cytokinesis. Molecular biology of the cell. 30:1791–1804.

Ng, A.Q.E., A.Y.E. Ng, and D. Zhang. 2020. Plasma membrane furrows control plasticity of ER-PM contacts. Cell Reports. 30:1434-1446.e1437.

Omasits, U., C.H. Ahrens, S. Müller, and B. Wollscheid. 2014. Protter: interactive protein feature visualization and integration with experimental proteomic data. Bioinformatics. 30:884–886.

Palmer, C.P., E. Aydar, and M.B. Djamgoz. 2005. A microbial TRP-like polycystic-kidney-disease-related ion channel gene. The Biochemical journal. 387:211–219.

Parnell, S.C., B.S. Magenheimer, R.L. Maser, T.S. Pavlov, M.A. Havens, M.L. Hastings, S.F. Jackson, C.J. Ward, K.R. Peterson, and A. Staruschenko. 2018. A mutation affecting polycystin-1 mediated heterotrimeric G-protein signaling causes PKD. Human molecular genetics. 27:3313–3324.

Pazour, G.J., J.T. San Agustin, J.A. Follit, J.L. Rosenbaum, and G.B. Witman. 2002. Polycystin-2 localizes to kidney cilia and the ciliary level is elevated in orpk mice with polycystic kidney disease. Current Biology. 12:R378–R380.

Rhind, N., Z. Chen, M. Yassour, D.A. Thompson, B.J. Haas, N. Habib, I. Wapinski, S. Roy, M.F. Lin, D.I. Heiman, S.K. Young, K. Furuya, Y. Guo, A. Pidoux, H.M. Chen, B. Robbertse, J.M. Goldberg, K. Aoki, E.H. Bayne, A.M. Berlin, C.A. Desjardins, E. Dobbs, L. Dukaj, L. Fan, M.G. FitzGerald, C. French, S. Gujja, K. Hansen, D. Keifenheim, J.Z. Levin, R.A. Mosher, C.A. Muller, J. Pfiffner, M. Priest, C. Russ, A. Smialowska, P. Swoboda, S.M. Sykes, M. Vaughn, S. Vengrova, R. Yoder, Q. Zeng, R. Allshire, D. Baulcombe, B.W. Birren, W. Brown, K. Ekwall, M. Kellis, J. Leatherwood, H. Levin, H. Margalit, R. Martienssen, C.A. Nieduszynski, J.W. Spatafora, N. Friedman, J.Z. Dalgaard, P. Baumann, H. Niki, A. Regev, and C. Nusbaum. 2011. Comparative functional genomics of the fission yeasts. Science (New York, N.Y). 332:930–936.

Sato, D., G. Hernandez-Hernandez, C. Matsumoto, S. Tajada, C.M. Moreno, R.E. Dixon, S. O’Dwyer, M.F. Navedo, J.S. Trimmer, C.E. Clancy, M.D. Binder, and L.F. Santana. 2019. A stochastic model of ion channel cluster formation in the plasma membrane. J Gen Physiol. 151:1116–1134.

Shen, P.S., X. Yang, P.G. DeCaen, X. Liu, D. Bulkley, D.E. Clapham, and E. Cao. 2016. The Structure of the Polycystic Kidney Disease Channel PKD2 in Lipid Nanodiscs. Cell. 167:763–773 e711.

Sinha, D., D. Ivan, E. Gibbs, M. Chetluru, J. Goss, and Q. Chen. 2022. Fission yeast polycystin Pkd2p promotes cell size expansion and antagonizes the Hippo-related SIN pathway. Journal of cell science. 135.

Su, Q., F. Hu, X. Ge, J. Lei, S. Yu, T. Wang, Q. Zhou, C. Mei, and Y. Shi. 2018. Structure of the human PKD1-PKD2 complex. Science. 361:eaat9819.

Su, X., M. Wu, G. Yao, W. El-Jouni, C. Luo, A. Tabari, and J. Zhou. 2015. Regulation of polycystin-1 ciliary trafficking by motifs at its C-terminus and polycystin-2 but not by cleavage at the GPS site. Journal of cell science. 128:4063–4073.

Thevenaz, P., U.E. Ruttimann, and M. Unser. 1998. A pyramid approach to subpixel registration based on intensity. IEEE transactions on image processing. 7:27–41.

Vještica, A., M. Marek, P.J. Nkosi, L. Merlini, G. Liu, M. Bérard, I. Billault-Chaumartin, and S.G. Martin. 2020. A toolbox of stable integration vectors in the fission yeast Schizosaccharomyces pombe. Journal of Cell Science. 133:jcs240754.

Wang, N., I.J. Lee, G. Rask, and J.Q. Wu. 2016. Roles of the TRAPP-II Complex and the Exocyst in Membrane Deposition during Fission Yeast Cytokinesis. PLoS biology. 14:e1002437.

Wu, G., V. D’Agati, Y. Cai, G. Markowitz, J.H. Park, D.M. Reynolds, Y. Maeda, T.C. Le, H. Hou Jr, and R. Kucherlapati. 1998. Somatic inactivation of Pkd2 results in polycystic kidney disease. Cell. 93:177–188.

Young, M.E., T.S. Karpova, B. Brugger, D.M. Moschenross, G.K. Wang, R. Schneiter, F.T. Wieland, and J.A. Cooper. 2002. The Sur7p family defines novel cortical domains in Saccharomyces cerevisiae, affects sphingolipid metabolism, and is involved in sporulation. Mol Cell Biol. 22:927–934.

Zhang, D., A. Vjestica, and S. Oliferenko. 2012. Plasma membrane tethering of the cortical ER necessitates its finely reticulated architecture. Current Biology. 22:2048–2052.

